# PRRC2 proteins regulate translation initiation by promoting leaky scanning

**DOI:** 10.1101/2022.11.11.516176

**Authors:** Jonathan Bohlen, Mykola Roiuk, Aurelio A. Teleman

**Affiliations:** German Cancer Research Center (DKFZ), 69120 Heidelberg, Germany; CellNetworks - Cluster of Excellence, Heidelberg University, Germany; Heidelberg University, 69120 Heidelberg, Germany; Heidelberg Biosciences International Graduate School (HBIGS), Germany; National Center for Tumor Diseases (NCT), partner site; Laboratory of Human Genetics of Infectious Diseases, Necker Branch, Institut National de la Santé et de la Recherche Médicale U1163, Paris, France; University of Paris, Imagine Institute, Paris, France

## Abstract

Roughly half of animal mRNAs contain upstream Open Reading Frames (uORFs). These uORFs represent an impediment to translation of the main ORF since ribosomes usually bind the mRNA cap at the 5’ end and then scan for ORFs in a 5’-to-3’ fashion. One way for ribosomes to bypass uORFs is via leaky scanning, whereby the ribosome disregards the uORF start codon. Hence leaky scanning is an important instance of post-transcriptional regulation that affects gene expression. Few molecular factors regulating or facilitating this process are known. Here we show that the PRRC2 proteins PRRC2A, PRRC2B and PRRC2C regulate translation initiation. We find that they bind eukaryotic translation initiation factors and preinitiation complexes, and are enriched on ribosomes translating mRNAs with uORFs. We find that PRRC2 proteins promote leaky scanning past translation start codons, thereby promoting translation of mRNAs containing uORFs. Since PRRC2 proteins have been associated with cancer, this provides a mechanistic starting point for understanding their physiological and pathophysiological roles.

## INTRODUCTION

Translation of mRNA into protein, the final step of gene expression, is tightly regulated both because it is energetically expensive, and because it directly results in changes in a cell’s proteome, thereby allowing cells to quickly react to environmental changes. Much of translation regulation occurs at the initiation step (1). Translation initiation starts with recruitment of the 43S pre-initiation ribosomal complex to the mRNA cap, followed by 5’ to 3’ scanning towards the main Open Reading Frame (ORF). While scanning, the 43S preinitiation complex searches for a start codon that is embedded in a favorable sequence context. In roughly half of mammalian mRNAs, however, this first start codon does not belong to the main ORF(mORF), but rather to an upstream ORF (uORF) (2). These uORFs thereby reduce translation of the main ORF by hijacking the scanning 43S ribosome. Two mechanisms exist to attenuate or circumvent this uORF-mediated repression: reinitiation and leaky scanning. In reinitiation, the ribosome translates the uORF, but after termination it regains an initiator tRNA, resumes scanning towards the main ORF, and then successfully translates the mORF that is further downstream (3). In contrast, in leaky scanning the ribosome bypasses the uORF start codon and continues scanning towards the mORF (4).

Some factors are known to affect leaky scanning. eIF1 (5), eIF1A (6) and eIF5 (7) all control start codon selection and initiation fidelity and hence modulate the rate of leaky scanning. The levels of eIF1 and eIF5 are controlled in negative feedback loops to finetune start codon selection and leaky scanning (8–10). More recently eIF4G2 (also called NAT1) has also been implicated in facilitating leaky scanning (11).

We investigate here the family of PRRC2 proteins PRRC2A, PRRC2B and PRRC2C, whose molecular functions are only starting to be studied. The PRRC2 proteins were detected as mRNA interacting proteins in several transcriptome-wide RNA interactome capture screens (12, 13). PRRC2A was shown to be an m^6^A reader which in the mouse brain controls oligodendroglial specification and myelination in part via stabilization of the Olig2 mRNA (14). All three PRRC2 proteins are present in stress granules and PRRC2C is required for efficient stress granule formation, while PRRC2A and B were not tested (15). Although their molecular functions are not completely understood, the PRRC2 proteins appear to have clinical relevance. Mutations in PRRC2A are associated with multiple cancers, including non-Hodgkin-Lymphoma (16), lung cancer (17), ER-positive breast cancer (18) and hepatocellular carcinoma, where they promote progression and immune infiltration (19). A PRRC2B-ALK gene fusion was found in an aggressive malignant neoplasm (20), and PRRC2C was found to have a different pattern of alternative splicing in non-small lung tumors compared to healthy lung (21). PRRC2A is also associated to rheumatoid arthritis (22).

In this study we discover that PRRC2 proteins are regulators of translation initiation that promote translation of uORFs-containing mRNAs by promoting leaky scanning.

## MATERIAL AND METHODS

### Cell lines and Culture Conditions

We cultured HeLa cells in DMEM +10% fetal bovine serum (FBS) +100 U/ml Penicillin/Streptomycin (Gibco 15140122). We sub-cultured the cells using Trypsin-EDTA. HeLa cells were tested negative for mycoplasma and authenticated using SNP typing.

### Generation of knockout cell lines

HeLa PRRC2A, PRRC2B, PRRC2C, and corresponding double and triple knock-outs were generated with the CRISPR-cas9 system. sgRNAs listed in Supp. Table 3 were cloned using sequences from the Brunello library (53) and cloned into pX459V2.0 via the Bpi1 site. Isogenic HeLa WT were transfected with a plasmid encoding sgRNAs using Lipofectamine 2000 according to manufacturer’s recommendations (Life Technologies #11668500). 24 hours post-transfection, cells were re-seeded into medium containing 1,5 µg/ml puromycin (Sigma-Aldrich #P9620). Three days after puromycin selection, cells were moved into normal medium (1x DMEM, 10% fetal bovine serum, 1% Penicillin/Streptomycin) and cultured till confluency. Single clone selection was performed with the use of multiple dilutions, by seeding cells in 96 well format. Single clones were selected, expanded and tested for the loss of protein of interest by immunoblotting. Double and triple knock-outs were generated by sequential knock-out of the proteins.

### siRNA mediated mRNA depletion

Cells were transfected with Lipofectamine RNAimax for siRNA mediated knockdowns. We seeded cells at 200.000 cells per 6-well, transfected them with 3 µl 10 µM siRNA (or siRNA Tetraplex) and 9 µl Lipofectamine reagent. We then re-seeded cells for experiments 72 hours post transfection. Sequences of siRNAs are provided in Suppl. Table 1.

### Ribosome Footprinting

Selective 40S and 80S ribosome footprinting was carried out as published in (27, 32) and as briefly below.

### Preparation of non-denaturing cell lysates

We seeded Hela cells at 1.5 million cells per 15 cm dish in 20 ml growth medium two days before cell harvest. For cell harvest, we poured off growth medium and washed cells quickly with ice-cold washing solution (1x PBS 10 mM MgCl2, 800 µM Cycloheximide). After quickly pouring off the washing solution, we added freshly prepared crosslinking solution (1x PBS, 10 mM MgCl_2_, 0.025% PFA, 0.5 mM DSP, 200 µM Cycloheximide) to the cells. We incubated cells with crosslinking solution for 15 minutes at room temperature while slowly rocking. We poured the crosslinking solution off and inactivated the remaining crosslinker for 5 minutes with ice-cold quenching solution (1x PBS, 10 mM MgCl_2_, 200 µM Cycloheximide, 300 mM Glycine). After removal of the quenching solution we added 150 µl of lysis buffer (0,25 M HEPES pH 7.5, 50 mM MgCl2, 1 M KCl, 5% NP40, 1000 μM Cycloheximide) to each 15 cm dish, resulting in 750µL of lysate. Lysis was carried out at 4°C. Then we scraped cells off the dish and collected the lysate. After brief vortexing, we clarified lysates by centrifugation at 20.000xg for 10 minutes at 4°C.

### Sucrose gradient centrifugation

We collected Supernatant and determined the approximate RNA concentration using a Nanodrop photo-spectrometer. We added 100 U of Ambion RNAse 1 per 120 µg of measured RNA. To obtain polysome profiles, we did not add RNAse. We incubated the Lysates for 5 minutes at 4°C and then loaded them onto 17.5%-50% sucrose gradients and centrifuged them for 5 hours at 35.000 rpm in a Beckman Ultracentrifuge in the SW40 rotor. Then we fractionated Gradients using a Biocomp Gradient Profiler system. We collected 40S and 80S fractions for immunoprecipitation and footprint isolation. We used 40S and 80S fractions corresponding to roughly one or two 15 cm dishes for direct extraction of RNA for total footprint sample and we used 40S and 80S fractions corresponding to roughly ten 15 cm dishes for immunoprecipitation of initiation factor bound ribosomes, to these fractions, we added NP40 to 1% final concentration.

### Immunoprecipitation

For immunoprecipitation, we bound antibodies to protein A or protein G magnetic dynabeads (Thermo) according to the manufacturer’s instructions. For PRRC2C selective ribosome footprinting, we used 40 µl antibody (Biomol #A303-315A) for each the 40S and 80S immunoprecipitation. We washed Beads three times and then added them to the cell lysates or 40S or 80S fractions. We incubated Solutions with beads for 2 hours or over-night, rotating at 4°C. Then we washed the beads three times with bead wash buffer (20 mM Tris pH 7.4, 10 mM MgCl2, 140 mM KCl, 1% NP40), including a change of reaction-vessel during the last wash. We increased the Bead volume to ~500µl with bead wash buffer. Then we subjected total footprint fractions and IPed fractions to crosslink removal and RNA extraction: we added 55 µl (1/9^th^ of volume) of crosslink-removal solution (10% SDS, 100 mM EDTA, 50 mM DTT), 600 µl Acid-Phenol Chloroform (Ambion) and incubated the mixture at 65°C, 1300 rpm shaking for 45 minutes. We then placed Tubes on ice for 5 minutes, spun for 5 min at 20.000 g and washed the supernatant once with acid-phenol chloroform and twice with chloroform. Then we precipitated RNA with Isopropanol and subjected it to library preparation (see below). We used the organic phase to isolate the precipitated or total proteins. We added 300 µl of ethanol, then 1,5 ml of isopropanol and incubatd the solutions at −20°C for 1 hour. We then sedimented the precipitated proteins by centrifugation at 20.000g for 20 minutes, washed them twice with 95% ethanol 0,3 M Guanidine HCl, dried and resuspended them in 1x Laemmli buffer.

### Deep-sequencing library preparation

After RNA extraction from total and IP-purified fractions, we determined the RNA quality and integrity on an Agilent Bioanalyzer using the total RNA Nano 6000 Chip. For size selection, we ran the RNA on 15% Urea-Polyacrylamide gels (Invitrogen) and excised fragments of size 25-35 nt (80S libraries) and 20-80 nt (40S libraries) using the Agilent small RNA ladder as a reference. We extracted RNA from the gel pieces by smashing the gels into small pieces with gel smasher tubes and extracting the RNA in 0.5 ml of 10 mM Tris pH 7 at 70°C for 10 minutes. We removed Gel pieces and precipitated RNA using isopropanol. We then dephosphorylated the Footprints using T4 PNK (NEB) for 2 hours at 37°C in PNK buffer without ATP. Then we again precipitated and purified the Footprints using isopropanol. For 40S footprints, we depleted contaminating 18S rRNA fragments as follows. We used prevalent 18S rRNA fragments from the first round of 40S footprinting to design complementary Biotin-TEG-DNA oligonucleotides (sequences listed in (27)), ordered from Sigma-Aldrich). We then hybridized 100ng of RNA footprints to a mixture (proportional to occurrence of the fragment) of these DNA oligos (in 40x molar excess) in (0.5M NaCl, 20mM Tris pH7.0, 1mM EDTA, 0.05% Tween20) by denaturing for 90 sec at 95°C and then annealing by reducing the temperature by −0.1°C/sec down to 37°C, then incubating 15min at 37°C. We pulled out Hybridized species using Streptavidin magnetic beads (NEB) by incubating at room temperature for 15 minutes, and purifiying the remaining RNA by isopropanol precipitation. We then assayed Footprints using an Agilent Bioanalyzer small RNA chip and Qubit smRNA kit. We used 25 ng or less of footprint RNA as input for library preparation with SMARTer smRNA-SeqKit for Illumina from Takara / Clontech Laboratories according to the manufacturer’s instructions. We sequenced Deep-sequencing libraries on the Illumina Next-Seq 550 system.

For RNA-seq libraries, we extracted total cell RNA using TRIzol and performed library preparation using the Illumina TruSeq Stranded library preparation kit. We also sequenced these RNA-seq libraries on the Illumina Next-Seq 550 system.

### Assessment of Co-translational assembly from selective 80S ribosome footprinting data

We analyzed our previously published selective 80S ribosome footprinting data (NCBI Geo GSE139391) to detect instances of co-translational assembly. For read processing and alignment we used our data analysis pipeline described in (27). For single transcript plots, we used custom scripts (available at https://github.com/aurelioteleman/Teleman-Lab) to count reads in the selective and total 80S samples for each individual transcript. Then, the ratio between selective and total 80S samples was calculated. A custom program was then used to identify co-translational assembly events, available at https://github.com/aurelioteleman/Teleman-Lab which fit the footprinting profile to a 1-step-function. This was done by selecting every position as a possible breakpoint, calculating the average of the data on each side of the breakpoint, and then calculating the sum-of-squares for distance to the average on each side of the breakpoint. The breakpoint position with the least sum of squares was selected as the optimal breakpoint for the step function. The difference in the average on the two sides of the breakpoint was then reported, yielding the ‘co-translational assembly score’.

### Preparation of cell lysates with RIPA buffer

Lysate preparation for conventional western blotting (without immunoprecipitation) was carried out as follows: Cells were seeded at 250.000 cells per 6-well. The next day, medium was removed and cells were briefly washed with DMEM. After removal of the medium, 200µl of RIPA buffer containing 20U Benzonase, protease inhibitors and phosphatase inihiibitors were added to each well, and cells were scraped off with a pipette tip. Lysates were collected and centrifuged for 10 minutes at 20.000g at 4°C. The protein concentration in the supernatant as measured by BCA. According to the protein concentration, samples were diluted to equal protein concentration and then 5x Laemmli buffer was added at 1/4^th^ the sample volume. Samples were incubated at 95°C for 5 minutes and then subjected to western blotting.

### Western blotting

We ran protein solutions from sucrose gradients, immunoprecipitations and lysates on SDS-PAGE gels and transferred to nitrocellulose membrane with 0.2 µm pore size. After Ponceau staining, we incubated the membranes in 5% skim milk PBST for 1 hour, briefly rinsed them with PBST and then incubated them in primary antibody solution (5% BSA PBST or 5% skim milk PBST) overnight at 4°C. We then washed the Membranes three times, 15 minutes each in PBST, incubated in secondary antibody solution (1:10000 in 5% skim milk PBST) for 1 hour at room temperature, and then washed them again three times for 15 minutes. Finally, we detected chemiluminescence using ECL reagents and imaged them with a Biorad ChemiDoc imaging system. We did not strip membranes. Antibodies used for immunoblotting are listed in Suppl. Table 2.

### Cell proliferation assay

For cell proliferation assays, we carried out siRNA mediated knockdowns out as described above. 72 hours after siRNA transduction, we re-seeded cells into 96 well plates at 1000 cells per well in 100 µl and 5 wells per condition. For each day of assay, we seeded one plate. The proliferation curve was performed by harvesting the first plate on the day of re-seeding, and one plate every successive day and proliferation was assessed by Cell Titer Glo according to the manufacturer’s instructions.

### Cloning

The Lamin B1 5’UTR firefly and renilla luciferase reporters and the Lamin B1 5’UTR stuORF reporter are from (54). aRaf and cRaf reporters are from (32). The reporters for different 5’UTR lengths are from (27). Renilla luciferase reporters with 5’UTRs of various genes were cloned by amplifying the 5’UTR of the gene of interest from HeLa cell cDNA and cloning it into the renilla luciferase reporter plasmid between the HindIII and Bsp119l sites. For some very short 5’UTRs, the sequence was inserted into the reporter plasmid via direct oligo cloning using the oligos indicated in Suppl. Table 4. DR1 uORF mutants were generated by site-directed mutagenesis PCR with the use of oligos listed in Suppl. Table 4, such that ATGs were mutated to TAG, near-cognate codons 1&2 into CCG and near-cognate codon 3 to GCG. The PCR products containing the mutations were cloned back into the backbone between HindIII and Bsp119I. To subclone uORF1 and uORF2 into the Lamin B1 5’ UTR, the LamB1 reporter containing a Kpn2I site from (55) was used. The DR1 region containing uORF1 and uORF2 was amplified with oligos containing Kpn2I and BshT1 sites and then cloned into the Kpn2I site of the LamB1 reporter. Reporters bearing the overlapping uORFs with different kozak strength were generated by oligo cloning between the Kpn2I and Bsp119I sites of the LamB1-Kpn2I reporter using the oligos listed in Suppl. Table 4. All constructs were verified by sequencing. All 5’UTR cloned for this manuscript are listed, including their transcript IDs and sequences, in Suppl. Table 5.

### Dual-luciferase translation reporter assay

We carried out Translation reporter assays after siRNA mediated knockdowns out as follows: We transfected Cells with siRNAs as described above. 72 hours after transfection, we re-seeded into a 96-well plate. We seeded HeLa control cells at 8.000 cells per well, and PRRC2A+B+C knockdown cells at 16.000 cells per well. We transfected Cells 16-20 hours after re-seeding using Lipofectamine 2000. We used 100 ng of renilla luciferase plasmid and 100 ng of firefly luciferase plasmid per well. 4 hours after transfection, we exchanged the medium.

### In-gradient formaldehyde crosslinking for translation initiation complex analysis

The method was carried out as described at (28). Briefly, 80% confluent 15 cm dish of HeLa wild type cells was lysed on ice with 120 ul of lysis buffer (10 mM Hepes pH 7.5, 62.5 mM KCl, 2.5 mM MgCl 2, 1 mM DTT, 1 % Triton X-100, 100 mg/ml cycloheximide, RiboLock, Protease and Phosphatase Inhibitor Cocktail). The lysate was clarified by centrifugation 14000 rpm 15 minutes at 4C, and loaded onto 7-30% linear sucrose gradient with progressively increasing concentration of formaldehyde. The gradient was prepared with the use of Biocomp Gradient Master, by mixing 30% sucrose (w/v) containing 0.05% formaldehyde with 7% sucrose prepared in following buffer: 10 mM Hepes pH 7.5, 62.5 mM KCl, 2.5 mM MgCl 2, 1mM DTT, 100 mg/ml cycloheximide. The samples were ultracentrifuged in SW40Ti rotor at 35000 rpm for 5 hours at 4C, following fractionation with use of a Biocomp Gradient Profiler system. Fraction 1-14 were run of PAAG and analyzed for protein distribution by western blot as described above.

## RESULTS

### Co-translational assembly occurs within and across eIF complexes

Proteins can bind nascent polypeptides that are in the process of being synthesized by a ribosome as soon as the nascent polypeptide is long enough to exit the ribosome channel (Fig. 1A). These interactions occur between nascent polypeptides and chaperones, as well as between nascent polypeptides and interacting proteins of a complex, leading to ‘co-translational chaperoning’ and ‘co-translational assembly’ of protein complexes, respectively (23–26). Co-translational assembly thereby results in a chain of physical links from the interacting protein to the nascent polypeptide, to the translating ribosome, to the mRNA coding for the nascent polypeptide. Hence co-translational assembly can be detected via selective ribosome footprinting by immunoprecipitating a protein of interest and looking for footprints on the mRNA of nascent polypeptides (24).

**Figure 1:**
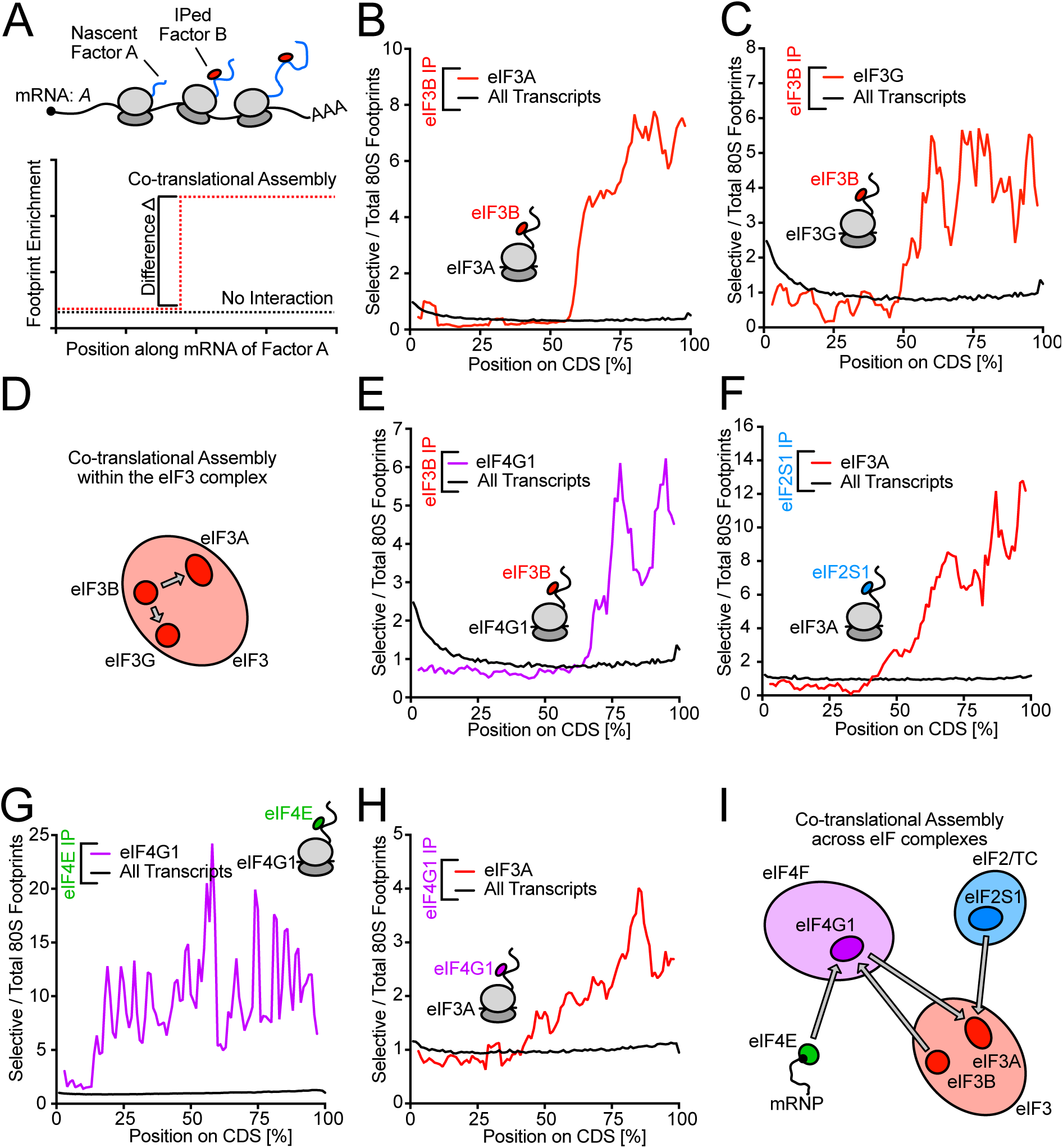
Selective ribosome footprinting reveals co-translational assembly within and across eIF complexes. **(A)** Concept of co-translational assembly and its detection by selective ribosome footprinting. In case of co-translational assembly, ribosomes bound by the bait factor X are pulled down due to interaction with the nascent polypeptide chain. Therefore, co-translational assembly will be visible as a sudden increase in the rate of factor X bound ribosomes observed on a given mRNA, with the position of this increase representing the onset of interaction between factor X and the nascent chain. **(B-D)** eIF3B interacts with nascent eIF3A (B) and eIF3G (C). Ratio of eIF3B selective 80S ribosome footprints per total 80S ribosome footprints on the eIF3A (B) or eIF3G (C) mRNA coding sequence (red) and on the main coding sequence of all other transcripts (black). Length of coding sequences is scaled between 0 and 100%. Data are an average of two biological replicates. (D) Summary of co-translational interactions within the eIF3 complex detected in eIF3B selective 80S ribosome footprinting. **(E-I)** Co-translational assembly between eIF complexes. (E) eIF3B interacts with nascent eIF4G1, (F) eIF2S1 interacts with nascent eIF3A, (G) eIF4E interacts with nascent eIF4G1 and (H) eIF4G1 interacts with nascent eIF3A. Data are an average of two biological replicates. (I) Summary of co-translational interactions between eIF complexes detected in selective 80S ribosome footprinting.

We previously published datasets from HeLa cells where we immunoprecipitated ribosomes bound to translation initiation factors (eIFs) of interest, such as eIF3B or eIF4G1, and then sequenced their footprints (27). To discover novel eIF interactors, we re-analyzed these data. We made use of the fact that footprints are detected on mRNAs encoding for interacting nascent polypeptides once the ribosome has finished translating the relevant interacting domain, causing a sudden increase in footprints on that mRNA (Figure 1A). Using an unbiased genome-wide analysis to search for such footprint profiles resembling a step-function, we detected co-translational assembly within the eIF3 complex whereby immunoprecipitation of eIF3B yielded footprints on the mRNAs of eIF3A and eIF3G (Figure 1B-D, Supplemental Data 1). This observation is consistent with a previous report of co-translational assembly of the eIF3 complex in yeast (28). We also detect co-translational assembly between different eIF complexes where we observe co-translational interaction of eIF3B with nascent eIF4G1, of eIF2S1 (eIF2α) with nascent eIF3A, of eIF4E with nascent eIF4G1 and of eIF4G1 with nascent eIF3A (Figures 1E-H). This suggests that eIF complexes start to engage in higher order assemblies already during translation of their subunits (Figure 1E-I).

### PRRC2 proteins assemble co-translationally with eIF3 and eIF4, and are associated to preinitiation complexes

Genome-wide, the top proteins that interact co-translationally with eIFs were other eIFs (Figure 2A-C). (Note that the protein being immunoprecipitated is often at the top of these lists because the antibody directly immunoprecipitates the nascent polypeptide once the epitope emerges from the ribosome). In addition, amongst the top interactors were the three members of the PRRC2 family: PRRC2A, PRRC2B and PRRC2C (Figure 2A-F, Suppl. Fig. 1A-B). Since these proteins have been linked to disease (16–19,22), we decided to study them in detail. We confirmed that PRRC2B and PRRC2C interact with eIFs by co-immunoprecipitation with endogenous eIF3B, eIF4G1 and eIF4E (Figure 2G). In the reverse setup, immunoprecipitation of PRRC2C led to co-precipitation of eIFs and of the ribosomal 40S component RPS15 (Figure 2H), suggesting it binds ribosomal preinitiation complexes. Knockdown of PRRC2A, PRRC2B and PRRC2C (Suppl. Figure 1C) caused the coIP signal to decrease (Figure 2H), confirming its specificity. On sucrose gradients, we observed that PRRC2B and PRRC2C sediment partly in polysomal fractions and upon RNase 1 treatment, they shift into 40S fractions (fractions 4-5, Suppl. Figure 1D), consistent with PRRC2 proteins interacting with pre-initiation complexes. Finally, a sedimentation protocol with in-gradient formaldehyde crosslinking that enables good resolution of pre-initiation complexes (29) also shows PRRC2C sedimentation in the 43-48S PIC peak (Suppl. Fig. 1E). Indeed, a larger fraction of total PRRC2C protein sediments in the PIC peak compared to eIF4G1 (compare fractions 12-14 versus 2-6, Suppl. Fig. 1E). Together with the fact that the copy numbers of PRRC2 proteins in HeLa cells are roughly in the same range as eukaryotic initiation factors (Suppl. Fig. 1F)(57), this suggests a large fraction of initiating ribosomes are bound by PRRC2 proteins. In summary these data indicate that PRRC2 proteins are interactors of eIFs and components of the translational machinery.

**Figure 2:**
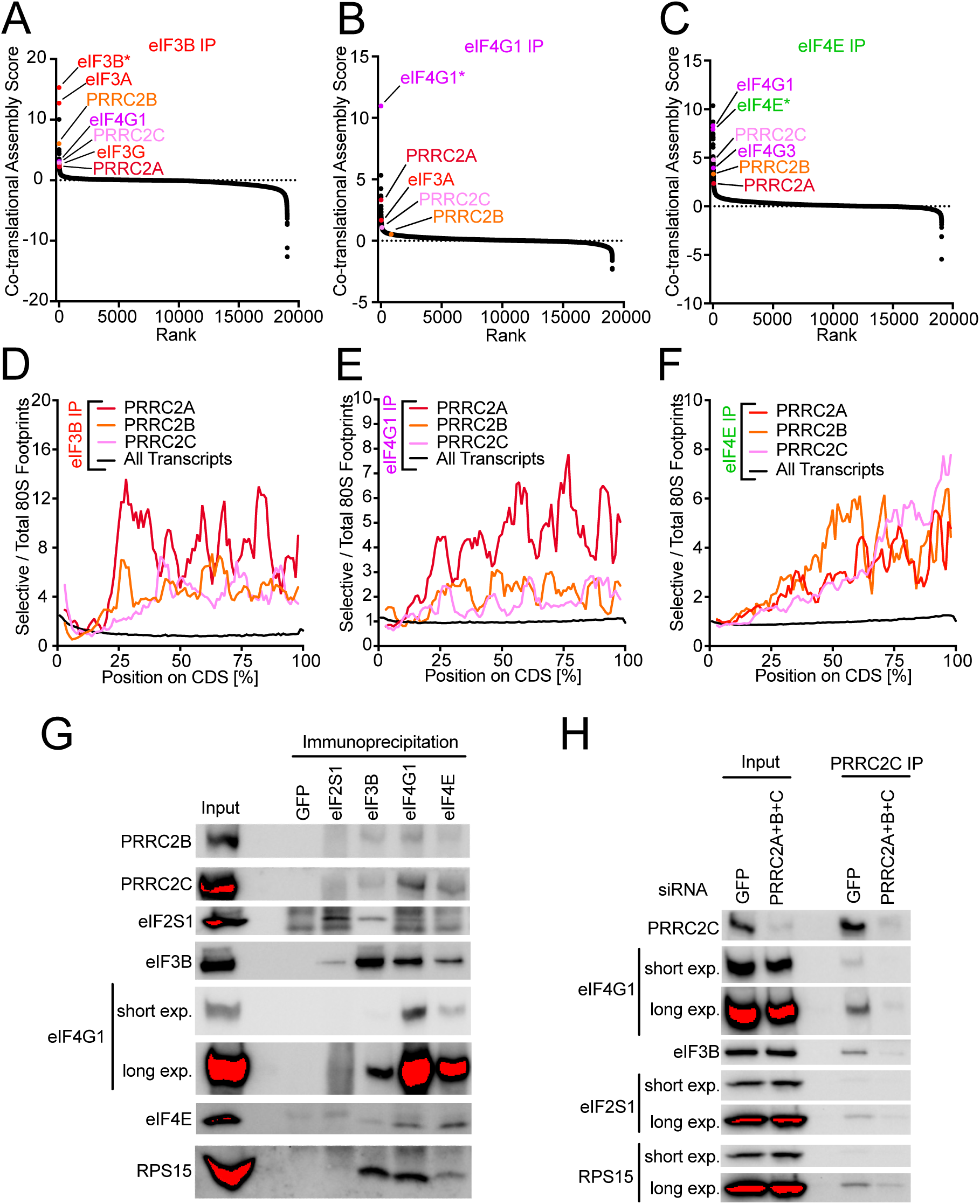
eIF3 and eIF4 interact co-translationally with PRRC2 proteins, which are associated to pre-initiation complexes. **(A-C)** PRRC proteins have high eIF3B **(A)**, eIFG1 **(B)** and eIF4E **(C)** co-translational assembly scores. Co-translational assembly scores, reflecting how well the footprint profile on an mRNA matches a step-function that shows increased binding as of a certain point on the profile, were determined as described in Methods, for all detected genes. Values were calculated from two biological replicates. **(D-F)** eIF3B, eIF4G1 and eIF4E interacts with nascent PRRC2A, PRRC2B and PRRC2C. Position-resolved number of eIF3B **(D)**, eIF4G1 **(E)** and eIF4E **(F)** selective 80S ribosome footprints normalized to total 80S ribosome footprints on the PRRC2A, PRRC2B and PRRC2C mRNA coding sequences and on all transcript coding sequence (black). Length of coding sequences is scaled between 0 and 100%. Data are an average of two biological replicates. **(G)** PRRC2B and PRRC2C co-immunoprecipitate with eIF3B, eIF4G1 and eIF4E. Immunoprecipitation of eIF3B, eIF4G1 and eIF4E, but not control IgG (anti-GFP) from whole-cell lysates of crosslinked HeLa cells co-precipitates PRRC2B and PRRC2C. Immunoprecipitations were done on cell lysates from cells crosslinked with formaldehyde and DSP as described in the Methods and (27). **(H)** Immunoprecipitation of PRRC2C co-precipitates pre-initiation complexes (PIC). Immunoprecipitation of PRRC2C from whole-cell lysates of crosslinked HeLa cells co-precipitates eIFs and ribosomal proteins. As a specificity control, siRNA mediated depletion of PRRC2A+B+C strongly reduces the amounts of co-precipitated PIC components. Immunoprecipitations were done on cell lysates from cells crosslinked with formaldehyde and DSP as described in the Methods and (27).

### PRRC2 family proteins are required for optimal proliferation and protein translation

To study the function of PRRC2 proteins in translation and cell proliferation we used siRNA-mediated RNA interference to deplete these proteins from HeLa cells. Co-depletion of PRRC2A and PRRC2C led to a significant increase in the cell doubling time (Fig. 3A-B, Suppl. Fig. 2A-D), which was further increased upon additional depletion of PRRC2B to almost twice the doubling time of control cells (Fig. 3A-B), indicating that HeLa cells require PRRC2 proteins for optimal proliferation. To confirm that this phenotype is on-target, we used two independent siRNAs targeting PRRC2C (siPRRC2C #1 and siPRRC2C #2). Both efficiently deplete PRRC2C protein in HeLa cells (Suppl. Figure 2E) and both cause an increase in cell doubling-time only when PRRC2A and PRRC2B are co-depleted (Suppl. Figure 2F). Finally, we also confirmed this phenotype using CRISPR/Cas9 mediated knockout of PRRC2A, PRRC2B and PRRC2C. Double-knockout cells lacking PRRC2A+C were significantly impaired in their proliferation (Fig. 3C-D). Although we were able to generate all the double-knockout combinations, we were not able to obtain complete triple-knockout cells, suggesting that complete loss of all PRRC2 proteins is lethal. Nonetheless, we obtained one line that was fully knockout for PRRC2B/C and had mutations in all 5 alleles of PRRC2A, albeit mainly triplet deletions leading to loss of >3 amino acids (Supplemental Figure 2G). This line (“PRRC2(A)BC KO”) has a strong proliferation impairment similar to the triple knockdown cells (Fig. 3C). Knockdown of all three PRRC2 proteins decreased protein translation rates, as determined by puromycin incorporation (Figure 3E) and polysome profiling (Figure 3F-G). In summary, PRRC2 proteins are required for optimal protein synthesis and cell proliferation in HeLa cells.

**Figure 3:**
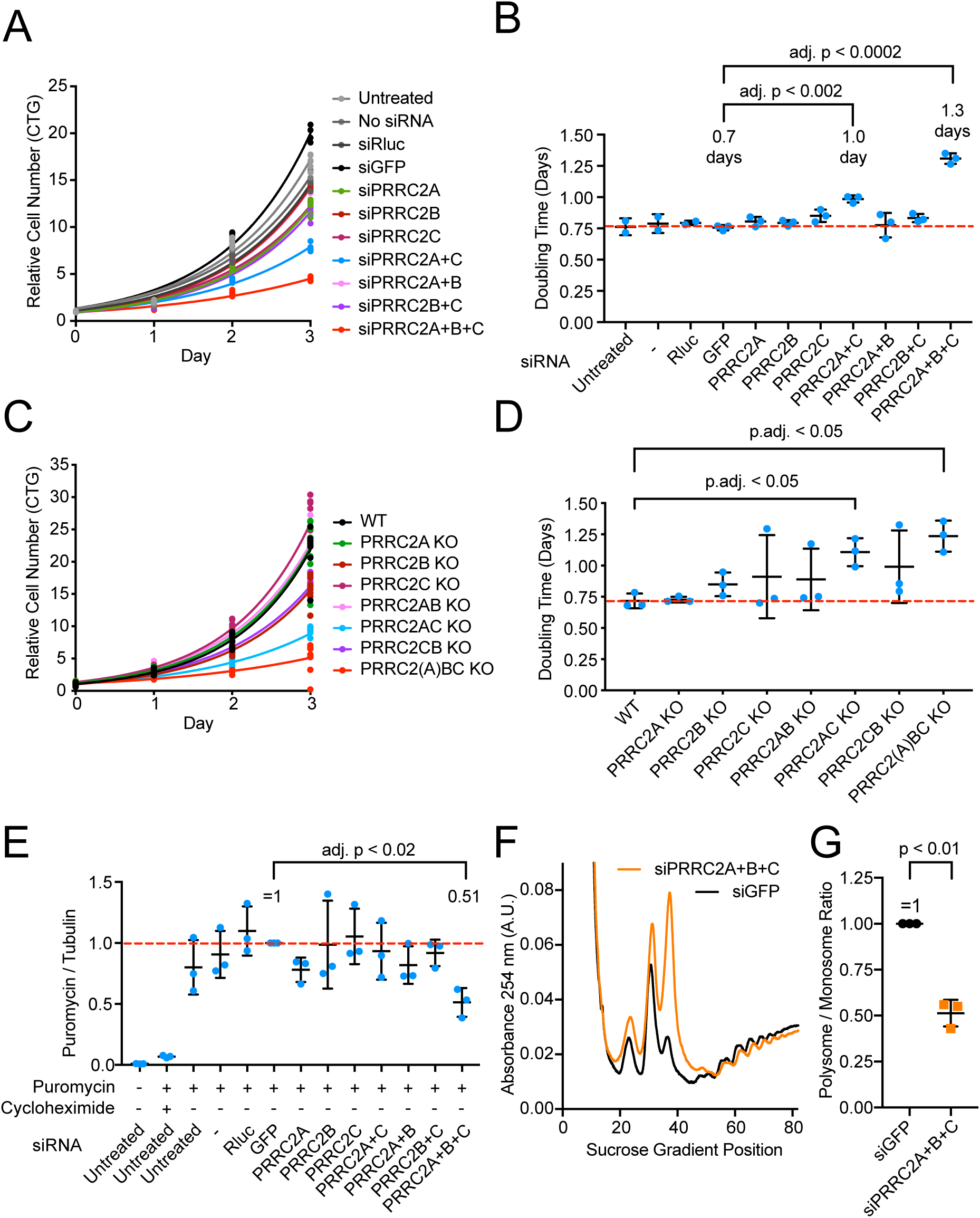
PRRC2 proteins promote proliferation and protein translation. **(A-B)** PRRC2 proteins are required for optimal cell proliferation. Proliferation of HeLa cells upon knockdown of PRRC2 genes was assayed using Cell titer glo (A) and doubling times were calculated by fitting the data to exponential functions (B). (A) representative experiment. siRNAs targeting Renilla luciferase (siRLuc) and GFP were used as negative controls. (B) Quantification of three biological replicates. P-values were calculated by multiple, two-sided t-tests, not assuming equal standard deviations and correcting for multiple testing. **(C-D)** Proliferation of HeLa cells lines knocked out for different PRRC2 proteins or combination of knockouts assayed using Cell titer glo. (C) Representative example. (D) Quantification of three biological replicates. **(E)** PRRC2 proteins are required for optimal protein translation. Protein synthesis in HeLa cells after PRRC2 protein depletion was assayed by Puromycin incorporation probed by western blotting. Data shown are from three biological replicates. P-values were calculated by multiple, two-sided t-tests, not assuming equal standard deviations and correcting for multiple testing. **(F-G)** PRRC2 proteins are required for optimal protein translation. Protein synthesis in HeLa cells after PRRC2 protein depletion was assayed by polysome profiling. (F) Representative polysome profile. (G) Quantification of the polysome/monosome ratio for three biological replicates shown. Statistical significant was assessed with paired, two-sided t-test, p = 0.0073.

We used publicly available ‘omic’ data to further investigate if PRRC2 proteins are essential. Both PRRC2B and PRRC2C whole-body knockout mice are not viable (30). PRRC2A KO mice have not been made, but PRRC2A KO in neurons causes significant impairment in neural development and survival (14). Additionally, heterozygous PRRC2C KO mice display abnormal phenotypes in behavior, vocalization, hearing and hair pattern (30). Consistent with these observations, predicted loss-of-function variants of PRRC2 genes are strongly depleted (10-fold less frequent than expected) from the general human population (31), which represents a strong signature of haploinsufficiency typical for eukaryotic translation initiation factors (eIF), ribosomal proteins (RP) and other essential genes. In summary, PRRC2 proteins are likely essential to mammalian life and subject to purifying selection to a similar degree as eIF and RP genes.

### PRRC2 family proteins are required for efficient translation of mRNAs containing uORFs

To study the role of PRRC2 proteins in mRNA translation, we performed 40S and 80S ribosome profiling (27), which detect footprints from initiating and elongating ribosomes, respectively (Figure 4A), in control cells or cells depleted for all three PRRC2 proteins (Figure 4B). On average, transcriptome-wide, we saw no obvious changes in the distribution of 40S footprints scanning in 5’UTRs (Suppl. Fig. 3A), or initiating on start codons (Suppl. Fig. 3B). In the absence of PRRC2A/B/C, there are fewer footprints aligning 25-40 nt upstream of start codons (Supp. Fig. 3B), which correspond to ribosomes directly on start codons but with large footprints due to the presence of initiation factors (27). Since this graph is normalized for the number of scanning 40S ribosomes approaching the start codon (80-100nt upstream), this potentially suggests that in the absence of PRRC2 proteins, ribosomes initiate more quickly at start codons. When we look at 80S footprints (Suppl. Fig. 3D-F), we observe a mild defect in termination or recycling, seen as an accumulation of 80S ribosomes on main ORF stop codons as well as the expected queuing of a second 80S ribosome 50nt in front of the stop codon (Suppl. Figure 3F). Unlike for DENR mutants (32) there is no defect in recycling of post-termination 40S ribosomes (Supplemental Figure 3C), suggesting that the recycling defect occurs at the level of termination or 80S splitting. Nonetheless, these global phenotypes are very mild, suggesting PRRC2 protein may rather play a role in translating specific mRNAs.

**Figure 4:**
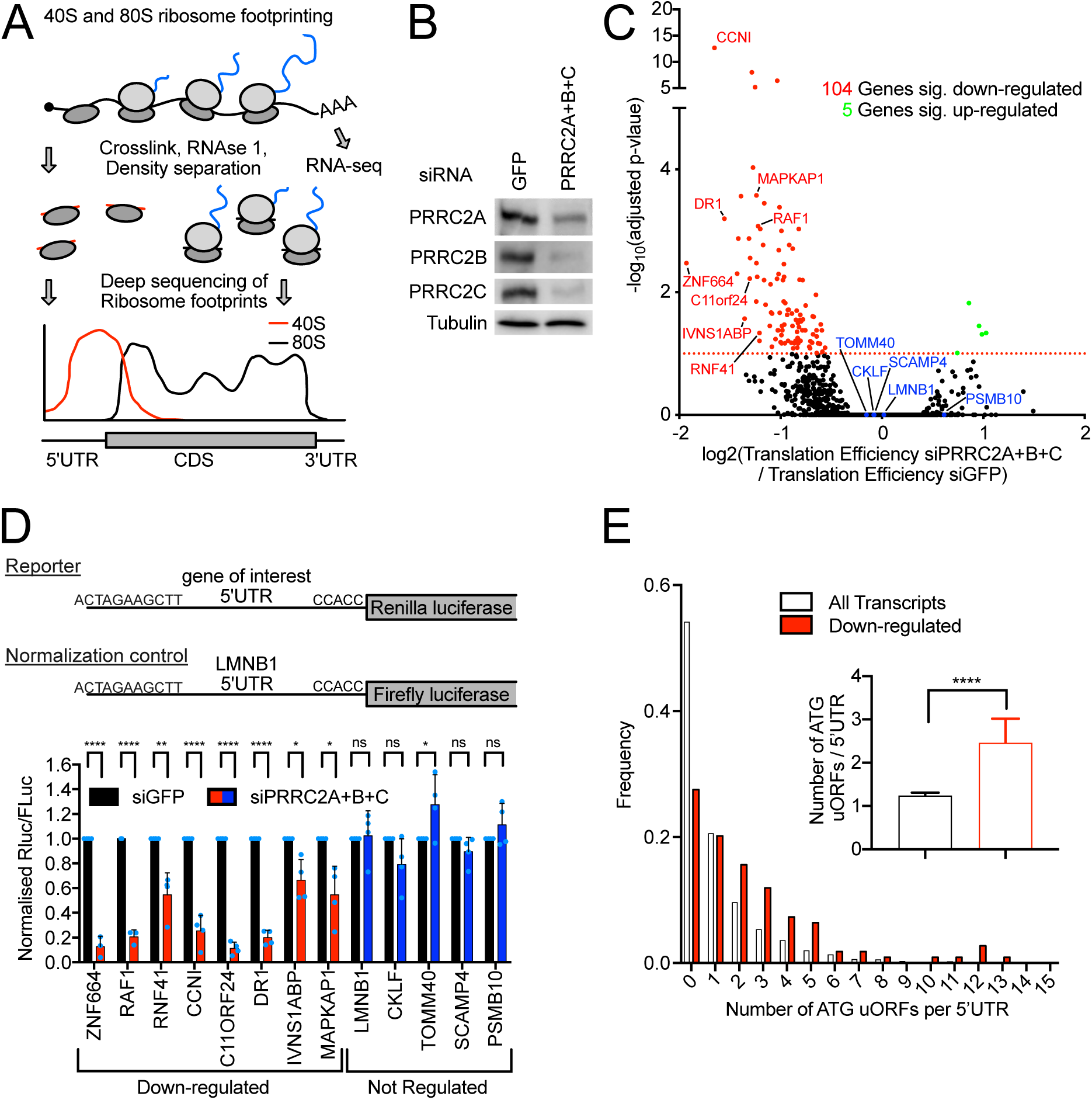
PRRC2 proteins promote translation of mRNAs containing uORFs. **(A)** Principle of 40S and 80S ribosome footprinting. Polysomes containing 40s scanning and 80S translating ribosomes are fixed in cells, RNAse treated in whole cells lysates and separated by density centrifugation. Ribosome footprints are prepared into libraries and analyzed by next-generation sequencing. **(B)** Efficiency of PRRC2 protein depletion for Ribosome footprinting experiments as determined by western blotting. Result is representative of two biological replicates. **(C)** PRRC2 proteins act as translational activators. Volcano plot showing 104 significantly down-regulated (red) and 5 significantly up-regulated (green) genes at the translational level upon PRRC2A+B+C depletion. Plot shows log_2_(fold-change in Translation Efficiency) and adjusted P-values for all detected genes, as obtained by X-tails analysis. **(D)** PRRC2 proteins regulate translation of mRNAs in virtue of their 5’UTRs. Activity of luciferase reporters carrying the 5’UTRs of down-regulated (red) and not-regulated (blue) genes from (C) upon PRRC2A+B+C depletion in HeLa cells. n=3-4 biological replicates. P-values calculated by unpaired, two-sided, t-test and adjusted for multiple testing. P-values: *p.adj.<0.05, **p.adj.<0.005, ****p.adj<0.00005, ns: p.adj.>0.05. **(E)** uORFs are enriched in PRRC2 regulated mRNAs. Histogram of the frequency of ATG-initiated uORFs per 5’UTR in PRRC2 dependent (n = 105) and independent (n = 5775) genes. Insert: Average number of ATG-initiated uORFs in PRRC2 dependent (n = 105) and independent (n = 5775) genes. Inset: mean number of uORFs per transcript; error bars: 95% confidence intervals. p-value < 0.0001 as calculated by Matt-Whitney test.

To test if there are specific mRNAs that are affected by PRRC2 depletion, we calculated translation efficiency (ribosome footprints / total mRNA) for each mRNA and found using Xtail (33) that mRNAs from 104 genes are translationally downregulated upon PRRC2 loss-of-function, whereas only 5 are significantly up-regulated (Figure 4C). Thus, PRRC2 proteins appear to promote translation, and to do so on a specific set of mRNAs. To confirm these results, we generated luciferase reporters carrying the 5’UTRs of eight predicted PRRC2 targets and five non-targets as negative controls. Indeed, knockdown of PRRC2A+B+C caused translation of the target reporters to drop, but not of the control reporters (Figure 4D). In addition to confirming the ribosome footprinting data, these results also indicate that the translation of these targets is PRRC2-dependent due at least in part to a feature in their 5’UTRs.

We noticed that PRRC2 targets have longer 5’UTRs compared to non-regulated mRNAs (Supplemental Figure 4A). To test if 5’UTR length *per se* is the factor determining PRRC2 dependence, we compared two luciferase reporters with 5’UTRs of differing length (26nt versus 728nt) whereby the longer 5’UTR was generated by multimerizing the shorter 26nt sequence, so as to increase its length without introducing additional regulatory elements (27). This shows that 5’UTR length does not determine PRRC2-dependence (Suppl. Fig. 4B), suggesting that PRRC2 targets carry specific features in their longer 5’UTRs.

We searched for features shared by 5’UTRs of PRRC2 targets and noticed that they are enriched for containing 2 or more uORFs (Figure 4E). Since this raises the possibility that uORFs cause PRRC2-dependence, we first asked if we could find a PRRC2-dependent gene that does not have uORFs in its 5’UTR. Of the 104 target genes, 16 do not have an ATG-initiated uORF. Since uORFs often start with a near-cognate start codon (34), we looked at our 80S footprinting data and found that 7 of these mRNAs had clear 80S footprints in their 5’UTRs, indicating the presence of actively translating ribosomes. Of the remaining 9 targets, 8 had clear 40S footprints in their 5’UTRs but not 80S footprints, identifying them as potential PRRC2 targets with no uORFs. We therefore cloned these 5’UTRs into luciferase reporters and tested their PRRC2-dependence. Unlike the luciferase reporters for the other target genes (Fig. 4D), none of these were PRRC2 dependent (Suppl. Fig. 4C), indicating that the selection for putative targets lacking uORFs strongly enriched for false-positives. In sum, all mRNAs whose translation we could confirm to be PRRC2-dependent, contain uORFs.

### uORFs are required for PRRC2-dependence

To test if uORFs are required for a 5’UTR to be PRR2C dependent, we analyzed the luciferase reporters carrying 5’UTRs of 3 PRRC2-targets – ARAF, RAF1 and DR1. Both ARAF and RAF1 have 2 uORFs that are clearly translated, as can be seen from the presence of 80S footprints in these regions (Fig. 5A-B). DR1 has a more complex 5’UTR with two uORFs that start with an AUG, and >7 putative uORFs that start with near-cognate start codons, some of which appear translated due to the presence of 80S footprints in these regions (Fig. 5C). As expected, all three reporters carrying the wildtype 5’UTR sequence of ARAF, RAF1 or DR1 are PRRC2-dependent (“wt”, Fig. 5D-F). We confirmed that these results are due to changes in protein translation and not mRNA levels by normalizing the Rluc activity to *Rluc* mRNA measured by qPCR and obtained the same results (Suppl. Fig. 4D). To test if the uORFs are required, we mutated the first two nucleotides of each start codon. For all three genes, combined mutation of the two uORF ATGs abolished or strongly diminished the PRRC2-dependence (Fig. 5D-F). Hence uORFs are a key element determining PRRC2-dependence. Consistent with this, introduction of a synthetic uORF composed of a start codon followed directly by a stop codon into a negative control luciferase reporter carrying the Lamin B1 5’UTR was sufficient to impart PRRC2-dependence (Fig. 5G, Suppl. Fig. 4E). Likewise, introduction of the two DR1 uORFs into the Lamin B1 reporter also rendered it PRRC2-dependent (Fig. 5G, Suppl. Fig. 4E). In sum, uORFs are required to render a 5’UTR dependent on PRRC2 for efficient translation.

**Figure 5:**
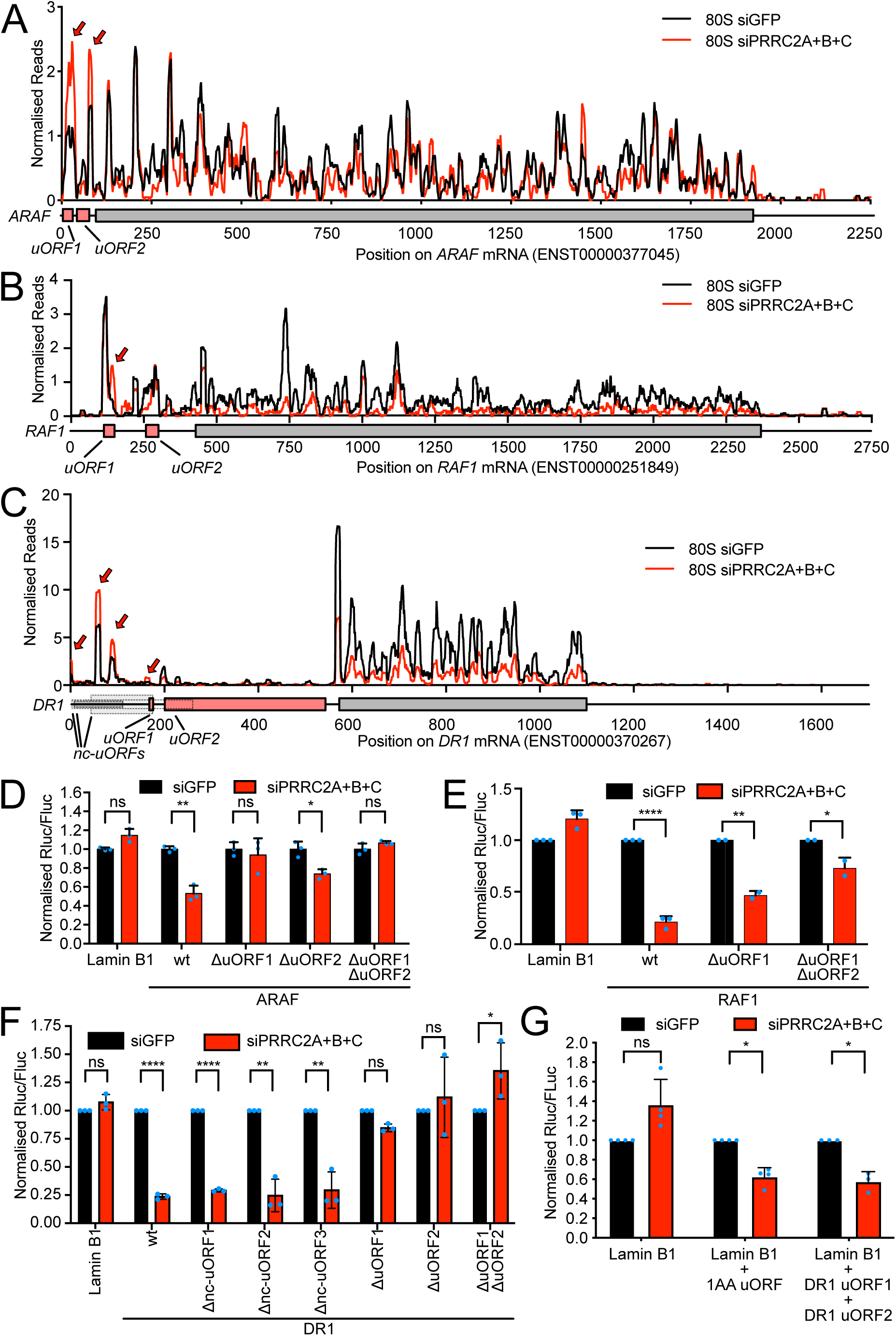
uORFs are required for PRRC2-dependence. **(A-C)** Ribosome footprints on ARAF (A), RAF1 (B) and DR1 (C) mRNAs in control (black) or PRRC2-knockdown cells (red). Read counts were normalized to sequencing depth. Graphs were smoothened with a sliding window of 11nt (A), 16 nt (B) or 10nt (C). mRNA features: uORFs (pink), main ORF (grey) and uORFs with near-cognate start codons (dotted lines) that have evidence for initiation from aggregate harringtonine traces in GWIPs-viz (56). Red arrow indicates 80S accumulation on uORF. **(D-F)** uORFs are required for PRRC2 dependence of ARAF(D), RAF1 (E) and DR1 (F) luciferase reporters. Mutation of the uORFs removes or severely blunts the PRRC2-depedence. n=2-3 biological replicates (E) or 3 biological replicates (D,F) +/- standard deviation. P-values calculated by unpaired, two-sided, t-test and adjusted for multiple testing. P-values: *p.adj.<0.05, **p.adj.<0.005, ****p.adj<0.00005, ns: p.adj.>0.05. **(G)** Presence of uORFs is sufficient to induce PRRC2 translational regulation. Dual-luciferase translation reporter assay of LMNB1 (negative control) 5’UTR reporter and LMNB1 reporter with insertion of 1AA uORF or DR1 uORF1 + uORF2. Schematic diagrams of the reporters are shown in Suppl. Fig. 4D. n= 3-4 biological replicates. P-values calculated by unpaired, two-sided, t-test and adjusted for multiple testing. P-values: *p.adj.<0.05, ns: p.adj.>0.05.

### PRRC2 proteins promote translation of uORF containing mRNAs via leaky scanning

The data presented above indicate that PRRC2 proteins somehow promote translation of a main ORF that is downstream of uORFs. There are essentially two mechanisms how ribosomes can overcome the inhibitory effect of uORFs. Either they scan past the uORF start codon in a process called ‘leaky scanning’, or they translate the uORF and then re-initiate translation downstream of the uORF (35). We noticed that upon knockdown of PRRC2A+B+C the number of 80S footprints within the uORFs of ARAF, RAF1 and DR1 mostly increase (red arrows, Fig. 5A-C). This suggests that in the absence of PRRC2 proteins, uORFs are more translated. Indeed, this trend can be observed transcriptome-wide. We previously identified a low-stringency set of all ‘translated uORFs’ as those that contain at least one 80S footprint (27). Metagene plots for 80S footprints relative to the start and stop codons of these translated uORFs shows that upon loss of PRRC2 proteins more ribosomes initiate translation and more ribosomes terminate translation on uORFs compared to control cells (Fig. 6A-B). Of note, genome-wide, the degree of accumulation of 80S ribosomes on uORF stop codons due to PRRC2 knockdown (Fig. 6B) is similar to the degree of 80S accumulation on uORF start codons (Fig. 6A) suggesting that the stop-codon accumulation reflects increased translation of uORFs and not a termination defect. These accumulations are not observed for scanning 40S footprints (Fig. 6C-D) indicating that in the absence of PRRC2, a higher proportion of the scanning 40S ribosomes convert into elongating 80S ribosomes on uORFs. Together, these data suggest that the presence of PRRC2 proteins helps promote leaky scanning.

**Figure 6:**
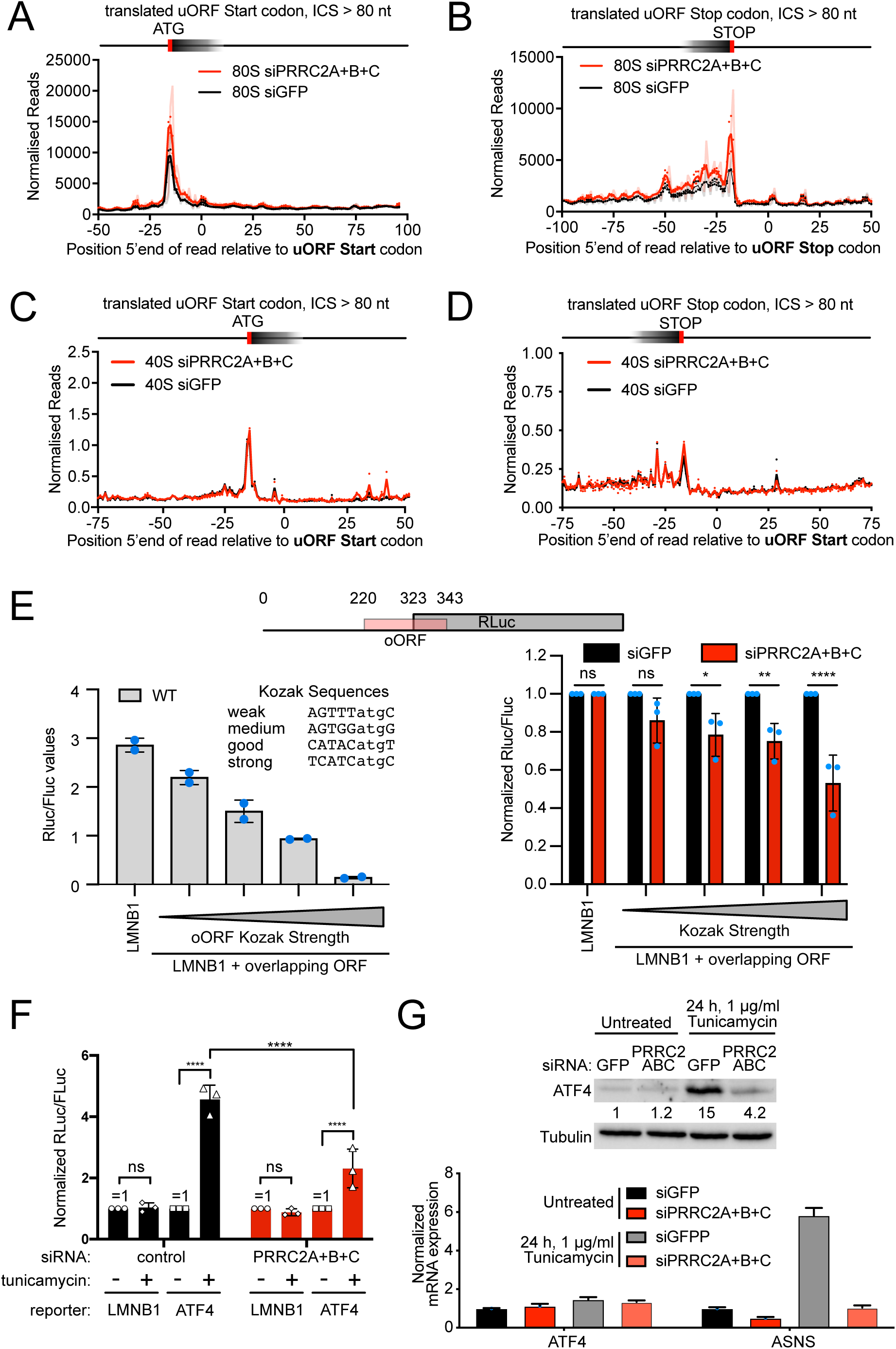
PRRC2 proteins promote leaky scanning. **(A-D)** PRRC2 knockdown causes accumulation of 80S ribosomes (A-B), but not scanning 40S ribosomes (C-D) on AUG-initiated uORFs. Metagene plots of 80S (A-B) or 40S (C-D) footprints near uORF start (A,C) or stop codons (B,D) of translated, AUG-initiated uORFs with an intercistronic space between uORF stop and mORF start codon of at least 80 nucleotides. Solid curves show data after triplet periodicity has been removed by averaging with a sliding window of 3 nt length. Shaded curves show non-smoothed data with triplet periodicity. **(E)** PRRC2 proteins promote leaky scanning. Luciferase reporter assay of Lamin B1 5’UTR reporters containing overlapping uORFs with start codons flanked by Kozak sequences of different strengths. Left panel: RLuc/FLuc ratios for the indicated reporters in wildtype cells shows the expected result that oORFs with stronger Kozak sequences inhibit RLuc translation more strongly. Right panel: PRRC2 knockdown reduces RLuc expression, and hence leaky scanning. n= 3 biological replicates +/- standard deviation. P-values calculated by unpaired, two-sided, t-test and adjusted for multiple testing. P-values: *p.adj.<0.05, **p.adj.<0.005, ****p.adj<0.00005, ns: p.adj.>0.05. **(F)** PRRC2 proteins are required for stress-dependent ATF4 induction via leaky scanning. Cells with PRRC2A+B+C or control siRNA knock-down were transfected with luciferase reporter bearing LMNB1 (negative control) 5’UTR or ATF4 5’UTR reporter. Induction of integrated stress response with 1ug/ml tunicamycin shows blunted induction of ATF4 reporter upon PRRC2A+B+C knock-down. n=3 biological replicates +/- standard deviation. P-values: *p.adj.<0.05, **p.adj.<0.005, ****p.adj<0.00005, ns: p.adj.>0.05 **(G)** PRRC2 knockdown causes impaired induction of ATF4 protein (top panel) and impaired transcriptional induction of the ATF4 target gene ASNS in response to stress (1µg/mL tunicamycin 24h).

To directly assay leaky scanning, we built a luciferase reporter containing an overlapping ORF (oORF) that overlaps with the luciferase coding sequence, extending 20nt past its start codon (Fig. 6E). Since ribosomes scan in a 5’-to-3’ direction, any ribosome that initiates on the oORF, and hence terminates downstream of the luciferase start codon, will not be able to translate luciferase. Thus, luciferase activity is a good readout for the number of ribosomes that (leaky) scan past the oORF start codon. The amount of leaky scanning depends on the strength of the oORF ATG sequence context. Hence, we constructed 4 oORF reporters with varying oORF ATG sequence context, from weak to strong as experimentally measured by (36). As expected, we observed that the stronger the oORF ATG context, the lower the signal of the luciferase mORF (Fig. 6E, left panel), thereby experimentally re-validating the strengths of the sequence contexts we selected. Knockdown of the PRRC2 proteins caused the luciferase signal to drop for all oORF constructs (Fig. 6E, right panel) indicating that PRRC2 proteins promote leaking scanning and not reinitiation. A similar effect was observed with PRRC2(A)BC KO cells (Suppl. Fig. 5A). The magnitude of the effect is larger when the oORF sequence context is strong, suggesting that PRRC2 proteins promote leaky scanning in particular when the sequence context is conducive to initiation. We hypothesized that the increased leaky scanning caused by loss of PRRC2 proteins should also occur on mORFs. It is challenging, however, to test this with luciferase reporters, because it would occur on the start codons of both the RLuc test reporter and the FLuc normalization control reporter, so this effect would be normalized away. Therefore, we made use of the fact that the knockdown of PRRC2 proteins reduced leaky scanning more strongly on start codons in a strong sequence context than a weak sequence context (Fig. 6E, right panel). We generated a series of reporters with differing ATG contexts on the RLuc main ORF (same sequences as in Fig. 6E), and co-transfected them with an FLuc normalization control with a medium ATG context. If loss of PRRC2 proteins increases recognition of mORF start codons in a strong sequence context more than in a weak sequence context, then the RLuc/FLuc ratio should increase as the RLuc sequence context gets stronger. Indeed, as expected, a stronger RLuc mORF ATG context led to a larger increase in signal upon PRRC2 knockdown, compared to weaker sequence context (Suppl. Fig. 5B), indicating that PRRC2 proteins also promote leaky scanning at mORF start codons. Furthermore, both experiments with the oORF or the mORF indicate that PRRC2 proteins preferentially decrease recognition of start codons in a strong sequence context.

One gene whose translation is regulated via a mechanism involving differential start-codon recognition is ATF4 (37–39). ATF4 has several uORFs that are translated, followed by an oORF that overlaps with the main ATF4 coding sequence and normally captures most of the ribosomes, causing ATF4 translation to be low. When the Integrated Stress Response is activated, scanning ribosomes bypass the oORF start codon and translate ATF4 instead. Hence, addition of tunicamycin to cells, which activates the Integrated Stress Response, causes an ATF4 luciferase reporter (32) to increase in expression (Fig. 6F). In agreement with PRRC2 promoting leaky scanning, induction of the ATF4 reporter is blunted upon PRRC2A+B+C knockdown (Fig. 6F). As expected,the overlapping uORF (uORF3) of ATF4 is required and sufficient to cause PRRC2-dependence (Suppl. Fig. 5C). Correspondingly, PRRC2 knockdown leads to strongly reduced levels of ATF4 protein, but not mRNA, in response to cell stress (Fig. 6G). In line with that observation, *ASNS* mRNA, a target of ATF4, is poorly induced in PRRC2A+B+C knockdown cells (Fig 6G, bottom panel). Thus, PRRC2 proteins are required for the proper induction of ATF4 upon cell stress. Since high tumor ATF4 activity strongly correlates with poor cancer prognosis (40–46), this could explain in part why PRRC2A is associated to cancer. Interestingly, since the mechanism by which the oORF in ATF4 is bypassed upon stress is thought to involve 40S ribosomes lacking an initiator tRNA, this suggests that PRRC2 proteins can promote leaky scanning by both ribosomes with a ternary complex, or ribosomes in the process of recruiting a new initiator-tRNA, for instance by increasing scanning speed.

Next, we tested whether PRRC2 depletion impairs leaky scanning by altering the levels of initiation factors involved in this process such as eIF1 or eIF5 (8–10). Levels of eIF5 or eIF1A are unchanged in cells depleted of PRRC2 by knockdown or knockout (Suppl. Fig. 6). Although eIF1 levels are elevated in PRRC2A+B+C KD cells (Suppl. Fig. 6A-B), they are unchanged in PRRC2(A)+B+C KO cells (Suppl. Fig. 6C-D), which also have elevated leaky scanning (Suppl. Fig. 5A). Thus the impaired leaky scanning observed in PRRC2 depletion conditions does not depend on altered levels of eIFs. Since PRRC2 proteins associate physically with PICs and eIFs, this is consistent with a more direct role of PRRC2 proteins in the translation initiation process.

### PRRC2C preferentially associate to ribosomes scanning and translating mRNAs with uORFs

We previously developed selective 40S footprinting as a method to detect when and where initiation factors bind to 40S ribosomes during the scanning and initiation process (27, 28). It works by immunoprecipitating 40S ribosomes bound to the protein of interest followed by sequencing of their footprints (Fig. 7A). To understand better how PRRC2 proteins function, we carried out selective 40S and 80S ribosome profiling for PRRC2C, since the PRRC2C antibody works for immunoprecipitation (Suppl. Fig. 7A). As no PRRC2C can be detected in the flow-through from both 40S and 80S precipitations, we conclude that the IP efficiently depletes the samples of PRRC2C-bound ribosomes. Metagene plots of these selective ribosome footprinting data (Suppl. Fig. 7B-G) revealed that on a transcriptome-wide level, PRRC2C binds 40S ribosomes evenly along 5’UTRs (Suppl. Fig. 7B). 80S ribosomes tend to be more highly bound by PRRC2C in 5’UTRs compared to coding sequences and binding of PRRC2 to 80S ribosomes decreases progressively during elongation (Suppl. Fig. 7E-F), as we previously observed for eIF3B, eIF4G1 and eIF4E (27). In sum, PRRC2C appears to behave similarly to standard eIFs, binding 40S ribosomes at cap recruitment and progressively dissociating from 80S ribosomes as they elongate.

**Figure 7:**
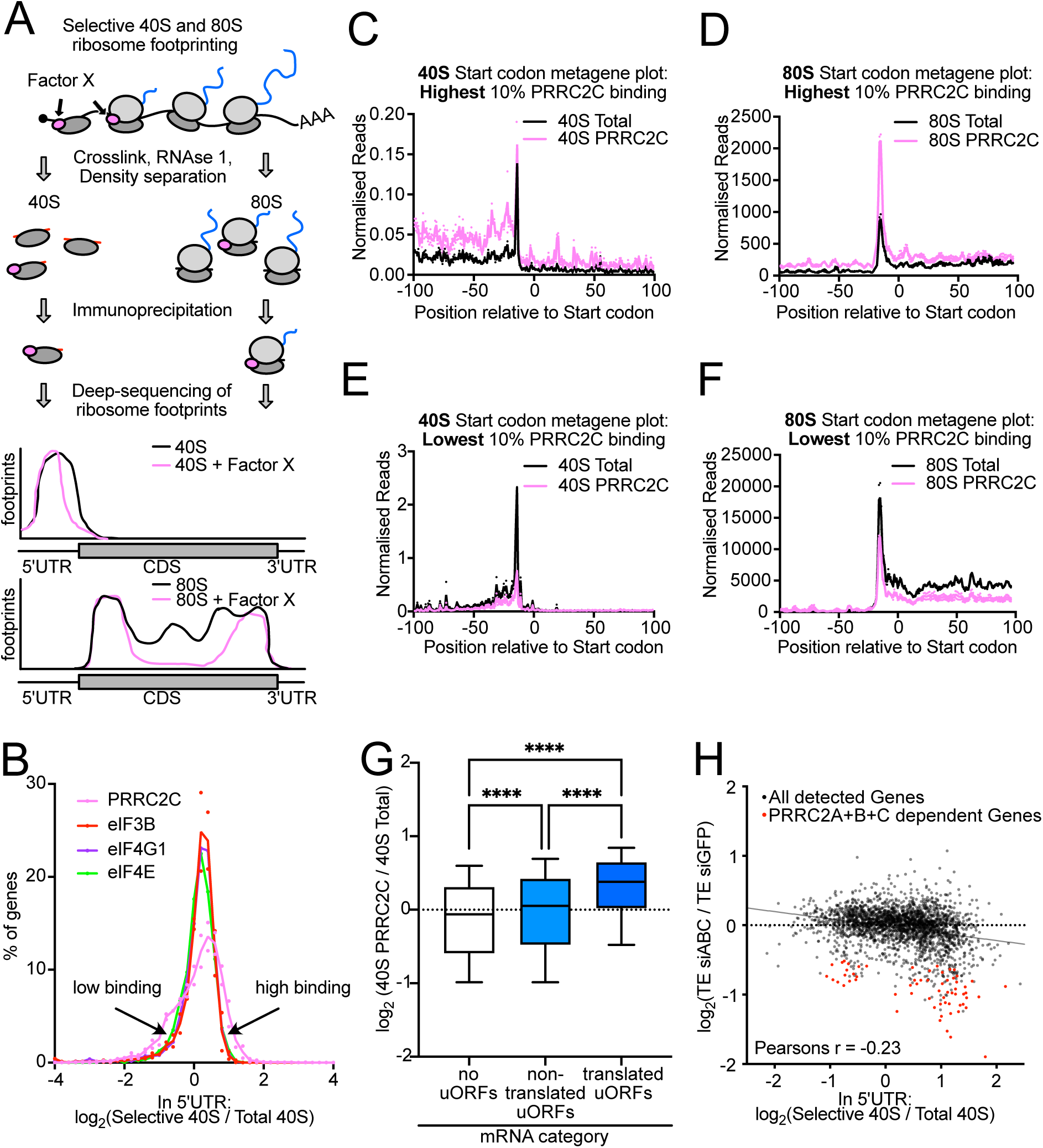
PRRC2C binding to mRNAs is heterogeneous and correlates to PRRC2-dependence. **(A)** Scheme and expected results of selective 40S and 80S ribosome footprinting. **(B)** PRRC2C binding to mRNAs is heterogenous when compared to binding of canonical eIFs. Histogram of the ratio of selective to total 40S ribosome footprints in 5’UTRs of all detected genes from PRRC2C, eIF3B, eIF4G1 and eIF4E selective 40S ribosome footprinting. n=2 biological replicates. **(C-F)** Footprints of ribosomes bound to PRRC2C are elevated throughout selected mRNAs. Start codon metagene plots of mRNAs with highest (top 10%, **C-D)** or lowest (bottom 10%, **E-F)** PRRC2C binding of 40S ribosomes in 5’UTRs. High binding to 40S ribosomes in 5’UTRs **(C, E)** predicts high binding to 80S ribosomes in 5’UTRs and coding sequences **(D, F)**. **(G)** Ribosomes containing PRRC2C are enriched on mRNAs containing translated uORFs. Transcripts were grouped into 3 categories: those without AUG-initiated uORFs (n=1967), those containing bioinformatically annotated uORFs but no detectable 80S footprints on those uORFs (“non-translated uORFs”, n=2003), and those with translated uORFs (n=2944). P values Kruskal-Wallis test followed by Dunn’s multiple comparison ****p.adj<0.0001 **(HH)** Genome-wide correlation between translational regulation by PRRC2 proteins and PRRC2C binding. Scatter plot of change in translational efficiency upon PRRC2A+B+C depletion versus PRRC2C binding rate to 40S ribosomes in 5’UTRs. Red dots: PRRC2-dependent transcripts from Fig. 4C. Plot shows the average of two biological replicates.

To ask whether PRRC2 binds equally to all mRNAs, we calculated for each detected gene (n=4707) the ratio of PRRC2C-selective 40S footprints to total 40S footprints. Interestingly, we found that PRRC2C displays heterogeneity in binding, with some mRNAs containing little PRRC2C binding and some a lot, in comparison to eIF3B, eIF4G1 and eIF4E, which bind roughly equally to all mRNAs (Figure 7B). This suggests that PRRC2C, as opposed to canonical translation initiation factors, has some specificity in its association to ribosomes on different mRNAs. To understand where this binding occurs on the mRNA, we grouped genes into two classes: the 10% with highest PRRC2 binding (n= 470) and the 10% with least binding (n = 470). mRNAs with high PRRC2 binding had high binding throughout – on scanning 40S in the 5’UTR and start codons (Fig. 7C) and on elongating 80S ribosomes (Fig. 7D) – whereas genes with low binding had low binding everywhere (Fig. 7E-F). The same pattern could be observed on single transcripts, such as RAF1, ARAF and DR1 (Suppl. Fig. 8). To test whether differential binding of PRRC2C to transcripts depends on uORFs, we grouped transcripts into 3 categories: those without ATG-initiated uORFs, those with uORFs but no detectable 80S footprints, suggesting they are not translated, and those with translated uORFs, as described above. This revealed that ribosomes containing PRRC2C are significantly enriched on transcripts with translated uORFs (Fig. 7G). Since we observe that high or low PRRC2C binding starts directly at the mRNA 5’cap (Suppl. Fig. 8D-E), it is not clear how the presence of a translated uORF further downstream in the 5’UTR regulates PRRC2C recruitment mechanistically (see Discussion below).

Finally, we asked whether mRNAs that have more PRRC2 binding depend more on PRRC2 for their efficient translation. Indeed, we observed a correlation between mRNAs with high PRRC2C binding and those that suffer a larger drop in translation upon knockdown of PRRC2A+B+C (Figure 7H). The drop in translation efficiency upon knockdown of PRRC2A+B+C for some mRNAs with low PRRC2 binding may be explained by PRRC2A or PRRC2B binding those mRNAs.

Thus, unlike the universal initiation factors eIF3B, eIF4G1 or eIF4E, we find that PRRC2C preferentially binds mRNAs containing translated uORFs, and that PRRC2 binding correlates with dependency on PRRC2 function for efficient translation.

## DISCUSSION

In this paper, we discover that PRRC2A, B and C are regulators of translation initiaton. They interact with eIFs and ribosomes, and are required for the efficient translation of mRNAs containing uORFs by promoting leaky scanning. As a consequence, PRRC2 protein depletion impairs mRNA translation and cell proliferation.

To discover these regulators of translation, we employed a new rationale by using selective ribosome profiling as a tool to discover new protein interactors. This works due to the fact that the protein of interest can interact with a nascent polypeptide already during its synthesis, co-precipitating the ribosome and allowing for the sequencing of the ribosome footprint on the mRNA of the nascent polypeptide. Previously, selective ribosome footprinting was used to study the action of co-translational chaperones (23–26) and translation initiation factors (27, 28). Therefore, this approach represents a new, sequencing-based method to discover protein-protein interactions that is complementary to existing methods. Particularly, the use of next-generation sequencing instead of protein mass-spectrometry may represent an advantage in terms of cost and sensitivity.

Leaky scanning is a mechanism of translational regulation first identified 30 years ago (47–49). Besides the core regulators of translation initiation (eIF1, eIF1A and eIF5), proteins influencing this process have remained largely elusive. Recently, eIF4G2/NAT1 was implicated in this process (11). We identify PRRC2 proteins as additional promoters of leaky scanning. PRRC2 proteins and NAT1 have been reported to interact physically (50), hence they may be working together. Since not all uORFs are subject to PRRC2 mediated regulation, there may be additional properties of PRRC2-dependent 5’UTRs that are yet to be uncovered (for example binding motifs in the vincinity of the uORF).

In addition to the defect in leaky scanning, we find that PRRC2 knockdown cells also have a termination or recycling defect on main ORF stop codons. Two observations suggest to us that this is likely an indirect consequence of PRRC2 knockdown, perhaps due to reduced translation of a factor involved in termination or recycling: 1) We find this defect is equally visible both on PRRC2 target mRNAs and non-target mRNAs. 2) We observe it equally on mRNAs bound highly by PRRC2C versus those bound weakly to PRRC2C. Therefore, we conclude that this is an indirect phenomenon, not directly linked to PRRC2 regulation or binding.

Previous work has linked PRRC2A to m6A modification of mRNA, describing PRRC2A as an m6A reader (14). Interestingly, m6A has also been linked to the leaky scanning mechanism regulating ATF4 induction (51). Hence it might be interesting in the future to study if there is a mechanistic link between m6A and leaky scanning. It will also be interesting in the future to study the biochemical function of PRRC2 proteins in *in vitro* translation systems, however it will require significant effort to obtain recombinant, soluble PRRC2 proteins since they are roughly 250-300 kD large.

PRRC2C displays a binding preference to a subset of mRNAs and may therefore mediate different rates of leaky scanning on different mRNAs in a cell. It is surprising that PRRC2C-high or low binding starts directly at the cap of the mRNA, suggesting that the degree of PRRC2C binding is determined and subsequently preserved at the recruitment of the PIC to the mRNA. How this works is unclear. There may be a mechanism in cis (e.g. by the presence of motifs anywhere in the mRNA) or in *trans* (by the subcellular microenvironment in which the mRNA is translated) that mediates the degree of PRRC2C binding.

PRRC2 proteins are among the few known translation factors that are not conserved from yeast to human. Instead, they are conserved among vertebrates (52), yet absent for example in *Drosophila melanogaster*, *Caenorhabditis elegans* and *Saccharomyces cerevisieae*. They may therefore provide a unique functionality to the translational machinery of these phyla. It is difficult to speculate why vertebrates may be more dependent on high rates of leaky scanning on subsets of their mRNAs. Answers might be provided by generation of KO strains or by the discovery of humans with mutations in PRRC2 proteins and characterization of their phenotype.

In sum, we identify here PRRC2 proteins as regulators of translation initiation that promote leaky scanning, enabling proteins encoded by ORFs downstream of uORFs to be more efficiently translated.

## DATA AVAILABILITY

All deep sequencing datasets have been submitted to NCBI Geo (GSE211440). Custom software is available on GitHub: https://github.com/aurelioteleman/Teleman-Lab.

## ACCESSION NUMBERS

All deep sequencing datasets have been submitted to NCBI Geo under accession number GSE211440.

## ACKNOWLEDGEMENTS

We thank Georg Stoecklin for critical reading of the manuscript and helpful comments.

## AUTHOR CONTRIBUTIONS

J.B. and M.R. performed experiments. All authors analyzed data and wrote the manuscript.

## CONFLICT OF INTEREST

Authors declare no competing interests.

## FIGURE LEGENDS

**Supplemental Figure 1:**
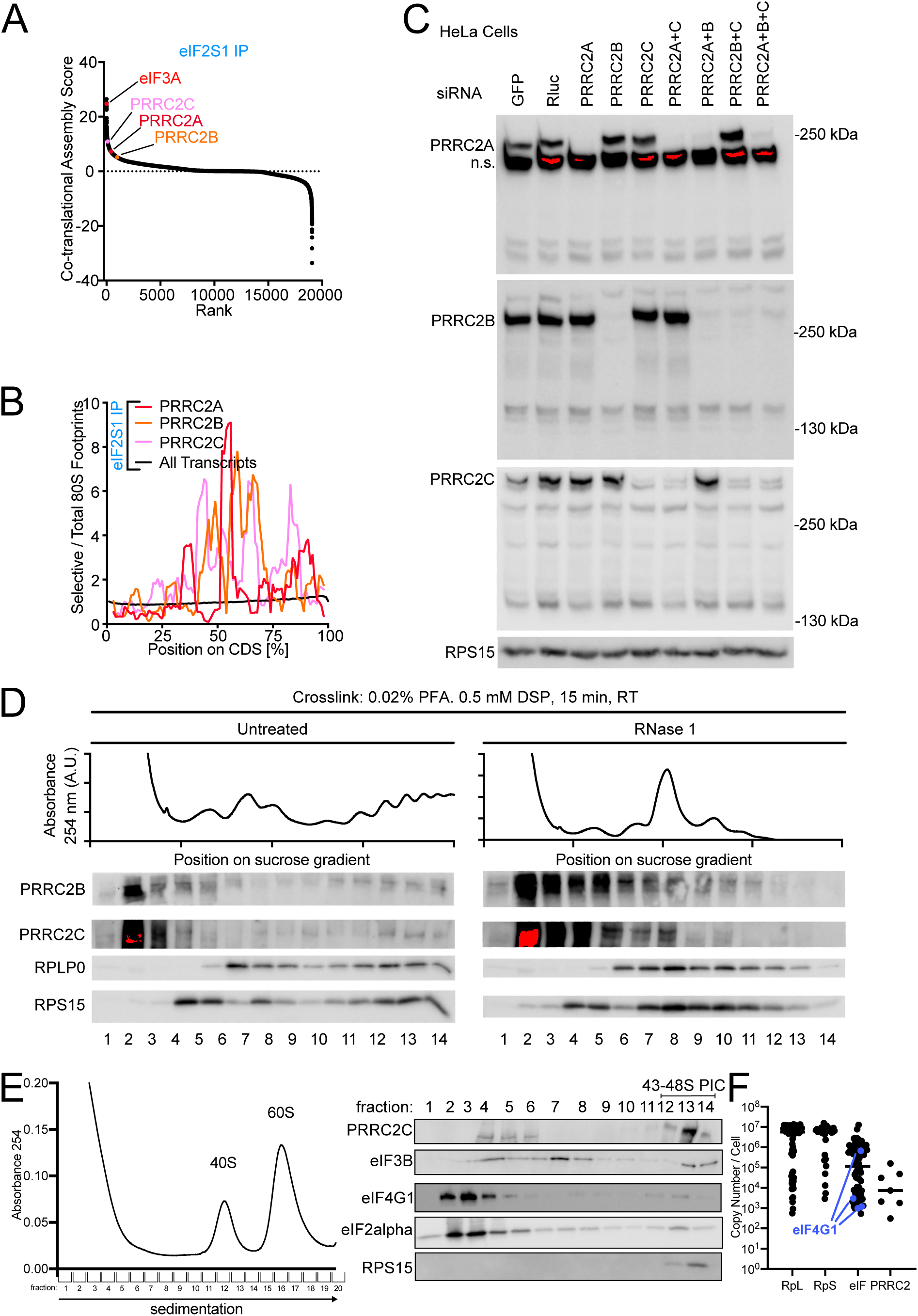
Co-translational interactions within and across eIF complexes. **(A)** PRRC proteins have intermediate to high co-translational assembly scores for interacting with eIF2S1. Co-translational assembly scores were determined as described in Methods for all detected genes. n=2 biological replicates. **(B)** eIF2S1 may interact with nascent PRRC2A, PRRC2B and PRRC2C. Ratio of eIF2S1 selective 80S ribosome footprints versus total 80S ribosome footprints on the PRRC2A, PRRC2B and PRRC2C mRNA coding sequences (colored) and on the main ORF of all other transcripts (black). Length of coding sequences is scaled between 0 and 100%. n=2 biological replicates. **(C)** PRRC2 protein depletion by siRNA transfection is efficient and specific. Knockdown efficiency and specificity of PRRC2 antibodies on lysates from HeLa cells treated with siRNA pools targeting the indicated genes. **(D)** PRRC2 proteins migrate with ribosomal and polysomal complexes in sucrose gradients. Polysome profiling and subsequent western blotting of gradient fractions from HeLa cells in the presence and absence of RNAse 1. RNAse 1 digestion causes polysomal PRRC2 proteins to run in 40S fractions 4-5. **(E)** Sucrose fractionation of translation initiation complexes. PRRC2C is detected in a free form (fractions 4-6) and co-sedimenting with translation initiation complexes (fractions12-14). **(F)** PRRC2 protein copy number in HeLa cells is roughly in the same range as the copy numbers of eukaryotic initiation factors (eIF). Data from (57) were re-analyzed. Each dot is one detected protein. Proteins were grouped into those of the large ribosomal subunit (“RpL”), small ribosomal subunit (“RpS”), initiation factors (“eIF”) or PRRC2 proteins.

**Supplemental Figure 2:**
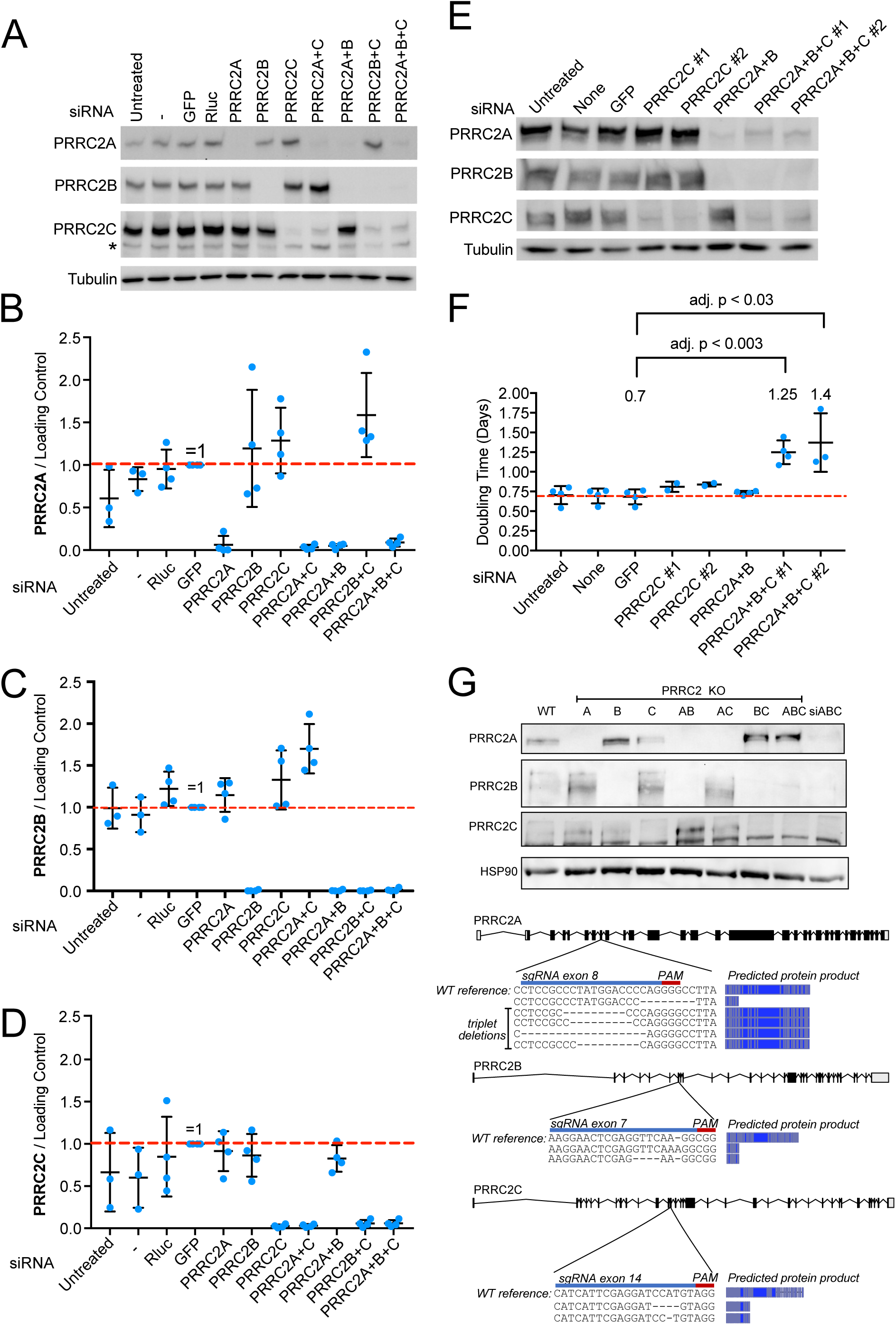
Knockdown of PRRC2 proteins causes reduced proliferation. **(A)** Immunoblot control of knockdown efficiency for the proliferation curves shown in main Figure 3A. **(B-D)** PRRC2 knockdowns are efficient and reproducible. Quantifications of PRRC2A **(B)**, PRRC2B **(C)**, and PRRC2C **(D)** knockdown efficiency in the indicated conditions as determined by western blotting. n=4 biological replicates. **(E)** Two independent siRNAs targeting PRRC2C efficiently knock down PRRC2C expression. Knockdown of PRRC2C with siRNA-1 and siRNA-2 in the presence or absence of PRRC2A and PRRC2B knockdown using siRNA pools, assayed by immunoblotting. n=3 biological replicates. **(F)** PRRC2C is required for optimal cell proliferation in the absence of PRRC2A and PRRC2B. Proliferation of HeLa cells after PRRC2 protein depletion was assayed using Cell Titer Glo and doubling times were calculated by fitting the data to exponential functions. n=3 biological replicates. P-values were calculated by multiple, two-sided t-tests, not assuming equal standard deviations and correcting for multiple testing. **(G)** Molecular characterization of the PRRC2(A)BC KO cell line. Both PRRC2B and C have mutations that cause frameshifts, resulting in truncated proteins, for all alleles detected. For PRRC2A, one allele has a frameshift mutation, whereas 4 alleles all have triplet deletions, leading to loss of >3 amino acids.

**Supplemental Figure 3:**
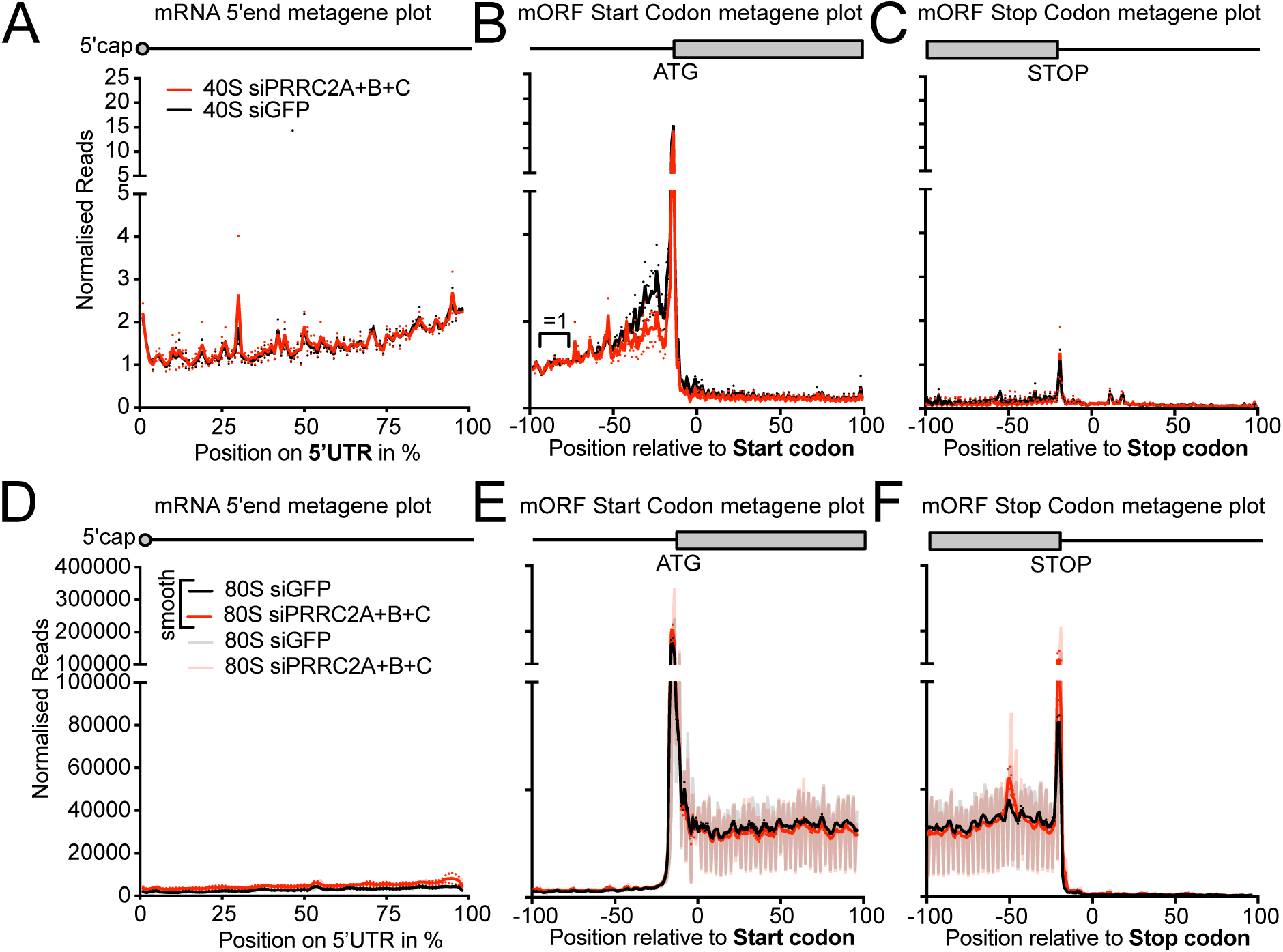
40S and 80S ribosome footprinting upon PRRC2 depletion. **(A-F)** Metagene plots of 40S **(A-C)** and 80S **(D-F)** ribosome footprints in control (black) or PRRC2 knockdown cells (red). Ribosome footprints were counted on all detected transcripts relative to the mRNA 5’UTR **(A, D,** scaled to 5’UTR length in %), main ORF start codon **(B, E)** or main ORF stop codons **(C, F)**. Solid curves in **(D-F)** show data after triplet periodicity has been removed by averaging with a sliding window of 3 nt length. Shaded curves show data without smoothing where triplet periodicity is visible.

**Supplemental Figure 4:**
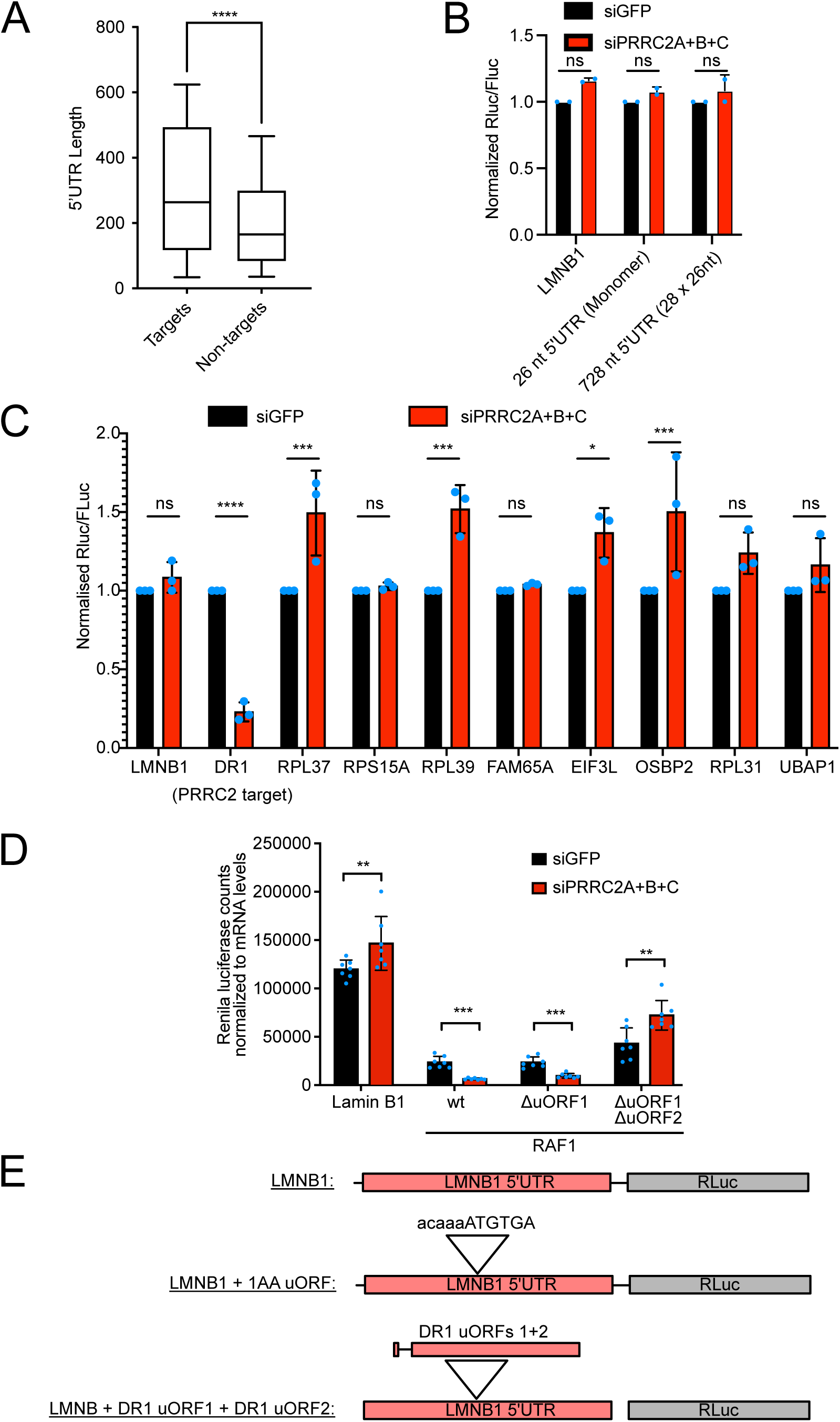
Presence of uORFs is required and sufficient for translational regulation by PRRC2 proteins via leaky scanning. **(A)** 5’UTR length of PRRC2 targets (as determined by X-tails analysis) and non-targets. Whiskers indicate top and bottom deciles while boxes indicate quartiles, the bar in the box represents the median. p-value < 0.0001 as calculated by Matt-Whitney test. **(B)** 5’UTR length does not affect PRRC2 dependence. Dual-luciferase translation reporter assay of RLuc reporters carrying the LMNB1 5’UTR (negative control), or a synthetic 5’UTR length composed of a 26nt sequence lacking uORFs, either singly or multimerized. Reporter activity assessed upon control or PRRC2A+B+C knockdown. n=2 biological replicates. **(C)** 5’UTRs lacking uORFs do not impart PRRC2 dependence to luciferase reporters. Dual-luciferase translation reporter assay of LMNB1 (negative control), DR1 (positive control) and uORF-less potential PRRC2 target 5’UTRs assessed upon control or PRRC2A+B+C knockdown. n=3 biological replicates. **(D)** Confirmation of translational regulation of the RAF1 5’UTR by PRRC2 proteins. The Rluc translation reporter signal for each 5’UTR construct was normalized to the respective Rluc mRNA levels determined by qPCR. P-values were calculated by unpaired, two-sided, t-test and adjusted for multiple testing. n=7 biological replicates. **(E)** Schematic diagram of the luciferase reporters used in main Figure 5G.

**Supplemental Figure 5:**
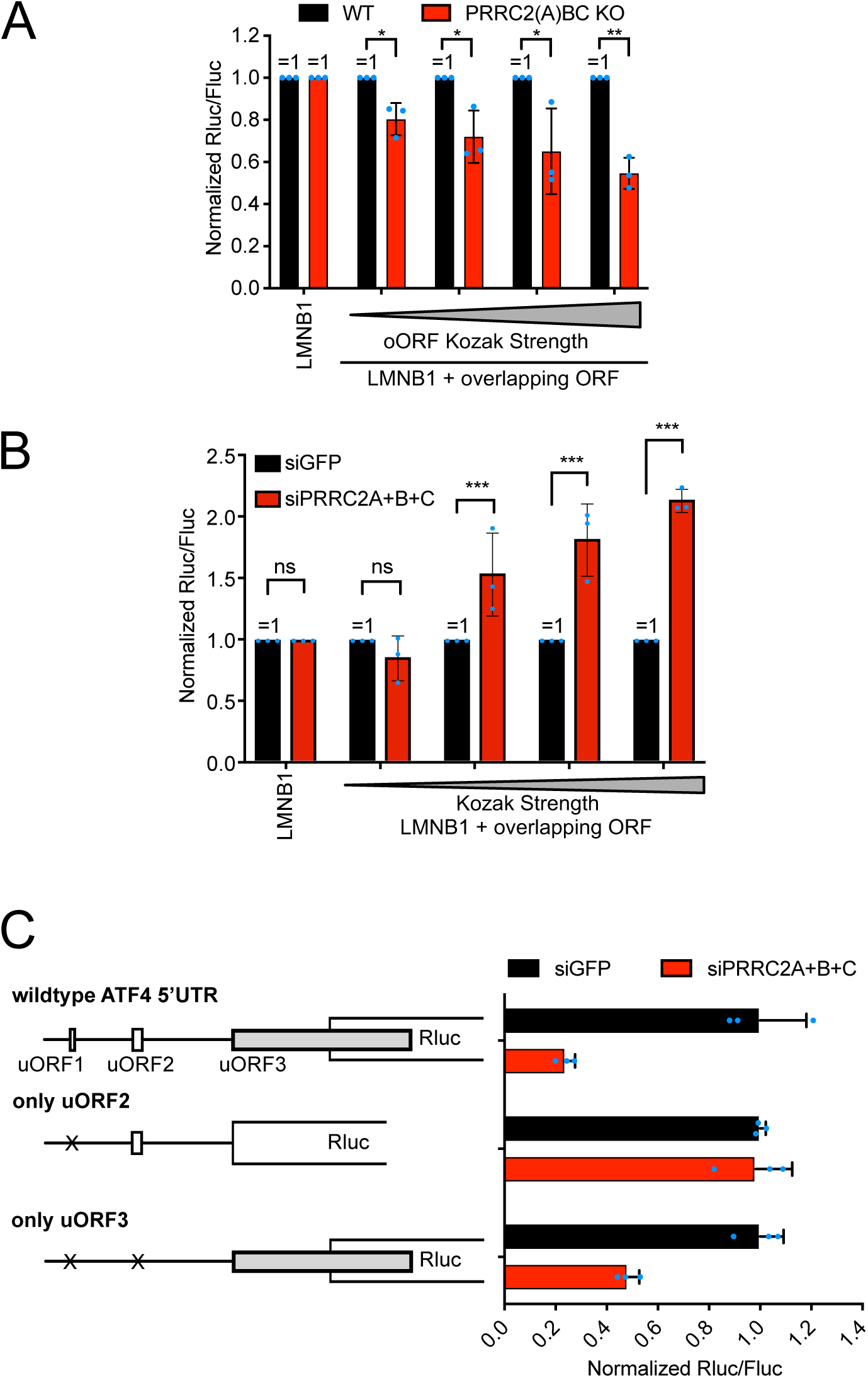
PRRC2 promotes leaky scanning. **(A)** PRRC2 proteins promote leaky scanning. Dual-luciferase translation reporter assay of a LMNB1 5’UTR reporter (negative control) and the same LMNB1 reporter with insertions of overlapping uORFs with start codons flanked by Kozak sequences of different strengths (as shown in main Fig. 6E). Activity assessed in control cells or PRRC2(A)BC KO cells. n=3 biological replicates. **(B)** Depletion of PRRC2 proteins causes increased leaky scanning also on the main ORF of Rluc. Dual-luciferase translation reporter assay on RLuc reporters with the LMNB1 (control) 5’UTR and start codon sequence contexts of different strengths (as shown in main Fig. 6E), normalized to an Fluc control reporter with the same 5’UTR (LMNB1). Activity assessed in control cells or PRRC2(A)BC KD cells. n=3 biological replicates. **(C)** The overlapping uORF3 of ATF4 is required and sufficient to cause a drop in expression of the ATF4-luciferase reporter upon PRRC2 knockdown.

**Supplemental Figure 6:**
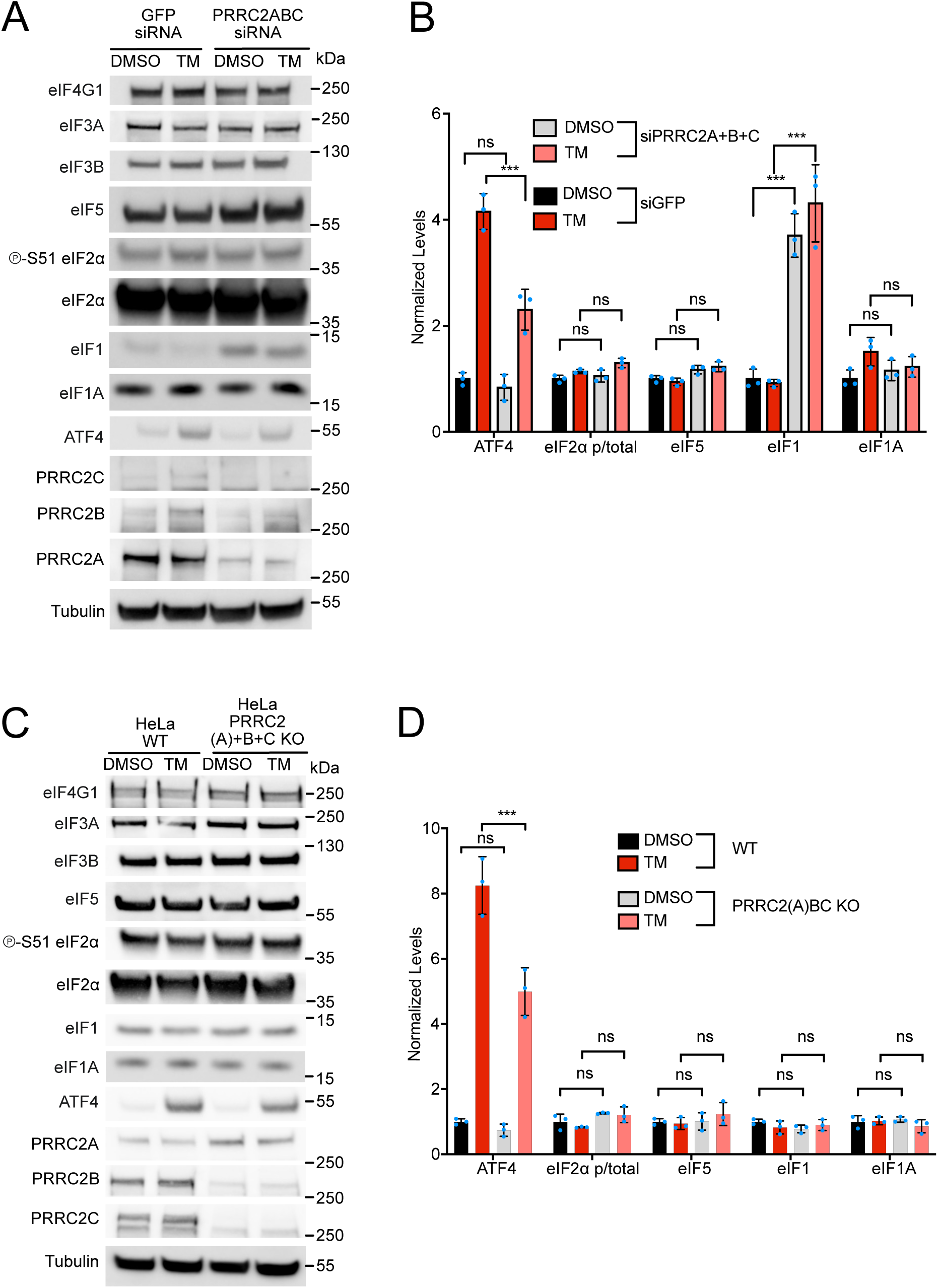
eIF levels in PRRC2 knockdown and knockout cells. HeLa cells depleted of PRRC2 proteins by the siRNA mediated (A-B) or sgRNA (C-D) were stimulated with tunicamycin for 16 hours and then harvested for western blotting. (A-B) PRRC2-protein depletion by siRNA mediated knockdown causes an increase in eIF1 and impaired induction of ATF4. (A) Representative western blot of three biological replicates quantified in (B). (C-D) PRRC2-protein depletion by CRISPR/Cas9 mediated knockout does not perturb eIF protein levels, but impairs ATF4 induction. (C) Representative western blot of three biological replicates quantified in (D).

**Supplemental Figure 7:**
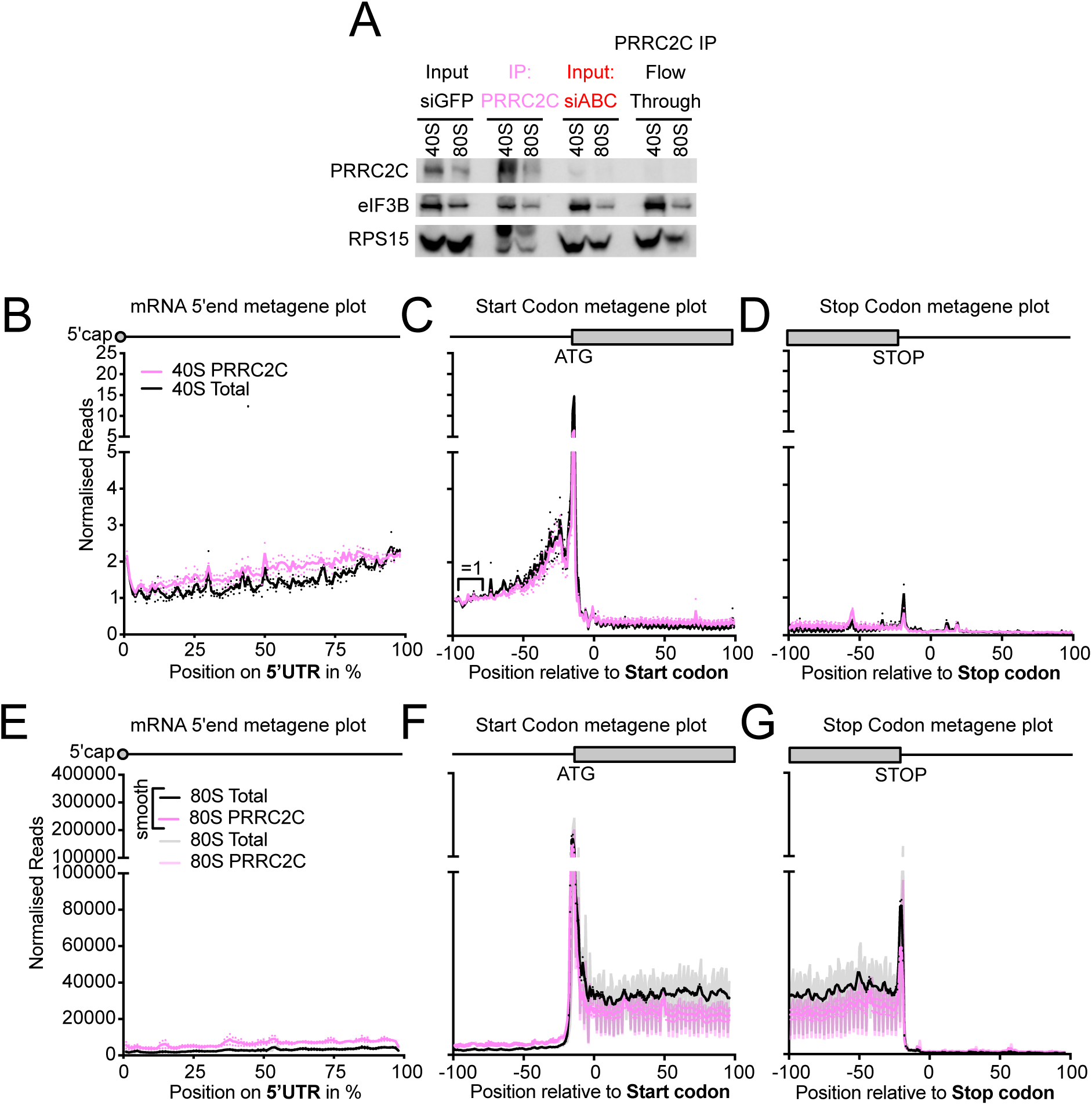
Quality controls for PRRC2C selective ribosome footprinting. **(A)** Western blot of input material, IPed material, and flow-through for control (siGFP) cells, or siPRRC2A+B+C knockdown cells, used for total or PRRC2C-selective ribosome footprinting. Representative of two biological replicates. **(B-G)** Metagene plots of total and PRRC2C selective 40S **(B-D)** and 80S **(E-G)** ribosome footprinting. Ribosome footprints were counted on all detected transcripts relative to mRNA 5’UTR **(B, E,** scaled to 5’UTR length in %), main ORF start codon **(C, F)** or main ORF stop codons **(D, G)**. Solid curves in **(E-G)** show data after triplet periodicity has been removed by averaging with a sliding window of 3 nt length. Shaded curves show data without smoothing, where triplet periodicity is visible.

**Supplemental Figure 8:**
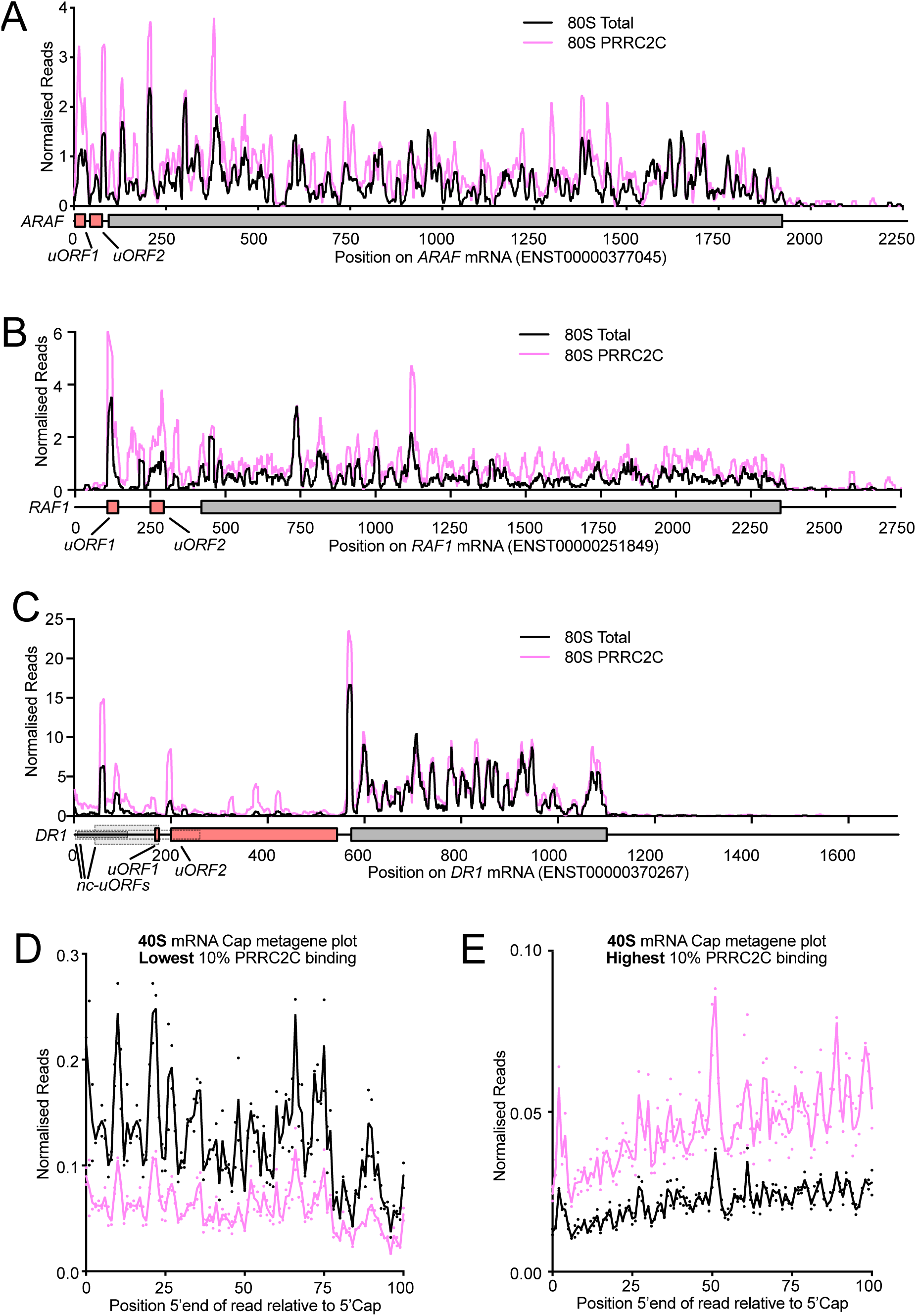
PRRC2C-selective ribosome footprints on target-gene mRNAs. **(A)** PRRC2C binding to 80S ribosomes on ARAF mRNA. Single transcript trace of total (black) and PRRC2C selective (pink) 80S ribosome footprints. Read counts were normalized to sequencing depth. Graphs were smoothened with a sliding window of 16 nt. mRNA features: uORFs (orange), mRNA (grey). Red arrow indicates 80S accumulation on uORF. **(B)** PRRC2C binding to 80S ribosomes on RAF1 mRNA. Single transcript trace of total (black) and PRRC2C selective (pink) 80S ribosome footprints. Read counts were normalized to sequencing depth. Graphs were smoothened with a sliding window of 16 nt. mRNA features: uORFs (orange), mRNA (grey). Red arrows indicate 80S accumulation on uORFs. **(C)** PRRC2C binding to 80S ribosomes on DR1 mRNA. Single transcript trace of total (black) and PRRC2C selective (pink) 80S ribosome footprints. Read counts were normalized to sequencing depth. Graphs were smoothened with a sliding window of 16 nt. mRNA features: uORFs (orange), mRNA (grey). Red arrows indicate 80S accumulation in the 5’UTR. **(D-E)** 5’Cap metagene plot of the 10% lowest (D) and highest (C) PRRC2C bound mRNAs. 40S Ribosome footprints were counted on all transcripts of the respective group relative to mRNA 5’Cap.

## Notes

### Competing Interest Statement

The authors have declared no competing interest.

